# The human RAD52 complex undergoes phase separation and facilitates bundling and end-to-end tethering of RAD51 presynaptic filaments

**DOI:** 10.1101/2024.12.09.627445

**Authors:** Ibraheem Alshareedah, Sushil Pangeni, Paul A. Dewan, Masayoshi Honda, Ting-Wei Liao, Maria Spies, Taekjip Ha

## Abstract

Human RAD52 is a prime target for synthetical lethality approaches to treat cancers with deficiency in homologous recombination. Among multiple cellular roles of RAD52, its functions in homologous recombination repair and protection of stalled replication forks appear to substitute those of the tumor suppressor protein BRCA2. However, the mechanistic details of how RAD52 can substitute BRCA2 functions are only beginning to emerge. RAD52 forms an undecameric ring that is enveloped by eleven ∼200 residue-long disordered regions, making it a highly multivalent and branched protein complex that potentiates supramolecular assembly. Here, we show that RAD52 exhibits homotypic phase separation capacity, and its condensates recruit key players in homologous recombination such as single-stranded (ss)DNA, RPA, and the RAD51 recombinase. Moreover, we show that RAD52 phase separation is regulated by its interaction partners such as ssDNA and RPA. Using fluorescence microscopy, we show that RAD52 can induce the formation of RAD51-ssDNA fibrillar structures. To probe the fine structure of these fibrils, we utilized single-molecule super-resolution imaging via DNA-PAINT and atomic force microscopy and showed that RAD51 fibrils are bundles of individual RAD51 nucleoprotein filaments. We further show that RAD52 induces end-to-end tethering of RAD51 nucleoprotein filaments. Overall, we demonstrate unique macromolecular organizational features of RAD52 that may underlie its various functions in the cell.

## Introduction

Genomic instability underlies many devastating diseases including different types of cancers and other DNA damage-related conditions^1-3^. Genomic instability usually arises from errors in repairing the genome after being exposed to damage^1^. Double-strand breaks (DSBs) are the most lethal type of DNA damage that can occur during DNA metabolism or as a result of reactive oxygen species (ROS) or exposure to irradiation^4^. DSBs are repaired in many ways in the cell, the most general of which are non-homologous end joining (NHEJ) and homologous recombination (HR)^5^. HR preserves genetic information by restoring the original DNA sequence using a homologous DNA sequence elsewhere in the nucleus^6^. HR occurs mostly in the G2/S phase where the presence of sister chromatid ensures the availability of DNA templates to be used for repair^7^. For HR-mediated repair of a DSB, the DNA ends are resected by a tri-protein complex known as the MRN complex^8^. Subsequently, long-range resection is achieved via Exo1 or DNA2, which leaves approximately 5kb-long single-stranded (ss)DNA overhangs at the two DSB ends^9,10^. These long ssDNA stretches are coated by RPA to protect them from nucleases^11,12^. Subsequently, RPA is replaced by RAD51 to form a RAD51-ssDNA nucleoprotein filament (NPF) called the presynaptic complex^13^. The RAD51 NPF conducts a homology search in the genome to capture a homologous dsDNA donor to be used as a template for the repair process^13,14^. Once the DNA template is found, the RAD51 NPF facilitates strand-invasion of the dsDNA donor, forming a D-loop that triggers the action of polymerases to synthesize DNA sequences based on the template^13^. Finally, the D-loop structure is resolved and the DNA is ligated, which completes the repair process^13^. In addition to its role in HR, the RAD51 NPF plays important roles in DNA replication, especially in protecting stalled and damaged replication forks from degredation^15^.

Many details and steps of HR are still under extensive investigation. The central step in HR is the assembly of the RAD51 NPF^14^, which requires the replacement of RPA at the ssDNA ends with RAD51, a process that depends on the action of mediator proteins^16^. The main mediators of RPA-to-RAD51 switching are BRCA2 in humans and Rad52 in yeast^17-19^. Interestingly, humans also have RAD52 which shares many biochemical features with its yeast counterpart, however, its mediator activity remains under debate. Some studies showed that human RAD52 does not have a clear mediator activity for the RAD51 NPF formation according to in vitro tests^17,20^. This is consistent with reports suggesting that RAD52 cannot fully compensate for BRCA2 deficiency in human cells^21^. In contrast, other studies showed stimulatory effects of RAD52 in D-loop formation^22,23^ and homology search^24^. This suggests that the role of RAD52 in HR may be context-dependent. It was long assumed that the human RAD52 is dispensable for the survival of human cells^20,25,26^. However, recent evidence highlighted a synthetic lethality relation between RAD52 and BRCA2, which indicated that RAD52 becomes critical for cell survival in conditions where BRCA2 is not functional^20,27-30^. Importantly, mutations in the BRCA2 gene are associated with cancers and genetic disorders such as Fanconi Anemia^3,31-34^. In sum, inhibition of RAD52 is lethal to cancer cells lacking BRCA2 but is not lethal to healthy cells with functioning BRCA2, which makes RAD52 a prime target for drug development in BRCA-related cancers. Consequently, understanding the function and behavior of RAD52 opens the door to developing cancer therapeutic strategies.

While the role of human RAD52 in HR remains unclear, studies have highlighted other functions of the protein. RAD52 possesses a single-stranded DNA annealing activity that is used by the cell to repair DSBs committed to homology-directed repair, but that cannot be repaired by HR^35,36^. In S-phase, RAD52 plays two important roles at stalled DNA replication forks, first protecting the fork from reversal by motor proteins^37^, and later cooperating with MUS81 nuclease to cleave the stalled forks^38^. RAD52 is also necessary to initiate mitotic DNA synthesis and is involved in telomeric DNA repair processes^39-41^. In telomerase-negative cancer cells, RAD52 is involved in the alternative lengthening of telomeres pathway^41^. Further research has shown that RAD52 has a high affinity towards G-quadruplex ssDNA structures and R-loops and can drive a reverse strand exchange reaction with both DNA and RNA^42-44^. In the oncological context, RAD52 forms foci upon DNA damage that are thought to be DNA repair centers as they colocalize with other DSB repair proteins such as RAD51^29,30,45-47^. RAD52 foci formation is also shown to be regulated by the cell cycle, with increased foci formation being associated with the G2/S phase, which is the same cell cycle phase where HR occurs^48^. Therefore, it is likely that when BRCA2 is dysfunctional, RAD52 plays a role in HR through mechanisms of self-assembly that result in foci formation. Despite renewed research interest in the biology and biochemistry of RAD52, the mesoscale self-assembly properties of the human RAD52 complex are still not well understood.

Crystal structure and cryo-EM studies have shown that the human RAD52 forms an undecameric ring or a decameric washer-like structure made of eleven or ten identical subunits^49-54^. The N-terminal half of the protein contains the oligomerization domain and two DNA-binding sites and can be wrapped by ssDNA^55^. The C-terminal domain of the protein is disordered and contains RPA and RAD51 binding sites^55-57^. This leads to a branched structure where the ring is surrounded by eleven 200 AA-long disordered regions, making it an example of a branched multivalent protein complex. We argue that such multivalency may lead to interesting consequences in the context of protein macromolecular assembly as it allows the RAD52 complex to bridge multiple molecules through its disordered C-terminal arm. Branched multivalency has been shown to confer unique condensation properties in other systems such as DNA nanostars^58^. This makes the self-assembly and heterotypic interactions of RAD52 relevant from a polymer physics perspective.

In this work, we characterize the supra-molecular self-assembly properties of RAD52 and its interactions with the RAD51 NPF and RPA. We first show that RAD52 undergoes phase separation at micromolar and nanomolar concentrations in aqueous and crowded environments, respectively. We further show that RAD52 condensates can recruit multiple key players in DSB repair such as RPA, ssDNA, and RAD51. Due to its protein and DNA binding sites, RAD52 phase separation can be regulated strongly by the presence of its interaction partners. We show that ssDNA facilitates RAD52 phase separation at low mixing ratios but suppresses the same at high mixing ratios, pointing toward a reentrant phase transition phenomenon^59^. In contrast, RPA enhances RAD52 phase separation in a monotonic fashion within our observation window. Moreover, we observe that RAD51-ssDNA complexes form fibrillar structures spanning tens of microns in length upon the addition of RAD52. This is true even in the presence of RPA, which is known to block RAD51-ssDNA interactions^60^. We hypothesized that these fibrillar structures are bundles of RAD51 nucleoprotein filaments. Indeed, single-molecule imaging reveals that RAD52 can induce clustering of individual RAD51 NPFs, thus creating the fibrils. Strikingly, we find that RAD52 can tether RAD51 NPFs, leading to the formation of exceptionally long filaments. These results are further validated with atomic force microscopy data. Collectively, our results show that the human RAD52 complex undergoes phase separation and induces clustering and end-to-end tethering of RAD51 nucleoprotein filaments.

## Results

### RAD52 undergoes homotypic phase separation

The human RAD52 protein forms an undecamer with a structured ring at the core that is enveloped by eleven disordered C-terminal regions (**Fig. 1 a&b**). Analysis of the linear net charge per residue shows that the entire RAD52 chain contains positively and negatively charged regions (**Fig. 1c**). Accordingly, we argued that phase separation may ensue at low salt concentrations via coulomb interactions due to attractive forces between oppositely charged segments. Thus, we prepared solutions of RAD52 at 30 mM NaCl concentration and varied the protein concentration from 0.1 µM to 5 µM. At no crowding conditions, the protein underwent phase separation at 5 µM concentration (**Fig. 1d&f and Fig. S1**). We then used a pseudo-inert polymer (PEG8000) as a molecular crowder to mimic the more crowded conditions in the cellular context. Adding 1 % wt/vol crowder reduced the condensation threshold to 1 µM (**Fig. 1d**). Further addition of PEG led to phase separation at concentrations as low as 250 nM (**Fig 1d**). This relatively low threshold for phase separation is attributed to the large size of the undecamer complex (∼10 nm ring diameter^49^) and the unusually high multivalency resulting from the eleven IDRs. Notably, the concentration of proteins and nucleic acids within the nucleus is predicted to be in the range of 100-200 mg/ml, which is equivalent to 10-20% crowding conditions^61^. To quantify the degree of phase separation, we used an image analysis approach that detects the droplet-dilute phase interfaces in the image using a Farid filter^62^ that computes the image derivative (**Fig. 1e**, see methods). The quantitative state diagram of PEG and RAD52 shows that RAD52 undergoes phase separation that is promoted by molecular crowding in a crowder concentration-dependent manner (**Fig. 1f**). We next probed the stability of RAD52 condensates against monovalent salts. The addition of monovalent salt (NaCl) suppressed phase separation, with the droplet dissolution occurring at ∼200 mM NaCl concentration (**Fig. 1g and Fig. S2a**). This indicates that the primary driving force of RAD52 phase separation is Columb interactions between the oppositely charged domains within the protein. Since in vitro recombination reactions usually require divalent salts and ATP hydrolysis, we tested whether RAD52 phase separation is affected by the presence of divalent salts as well as ATP. Our results show that at no-crowder conditions, neither calcium nor ATP significantly alters the RAD52 condensation capacity in the range of co-solute concentrations usually employed for in vitro recombination assays (1-20 mM) (**Fig. 1 h&i and Fig. S2 b&c**).

**Figure 1.**
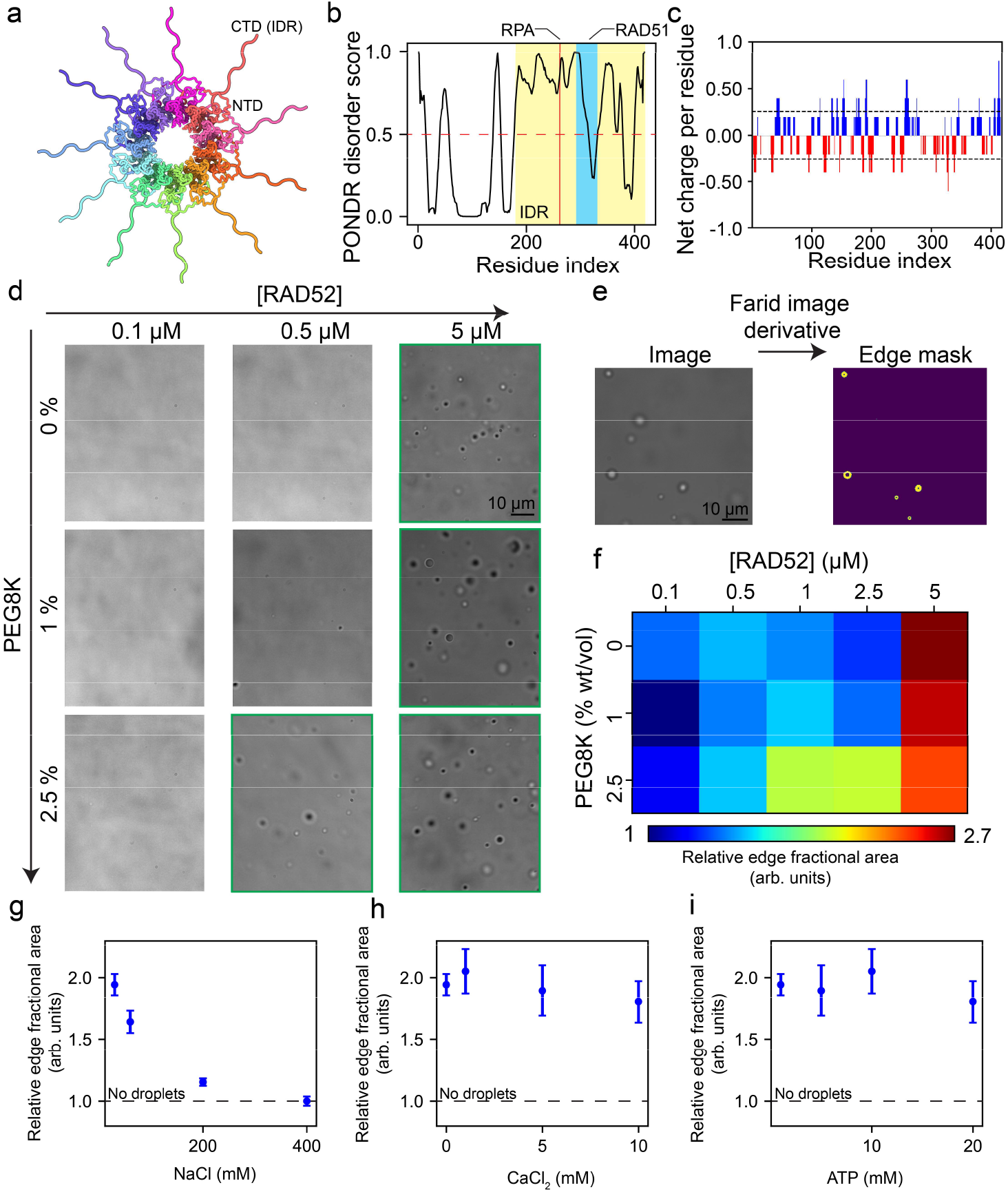
RAD52 undergoes homotypic phase separation. **(a)** A scheme showing the structure of the RAD52 undecameric ring visualized using Protein Imager^63^ based on a previously reported Cryo-EM structure^64^ (PDB ID 8BJM). The IDRs are manually drawn to illustrate their presence and are not to scale. **(b)** PONDR (predictor of natural disordered regions) score of the RAD52 protein showing that its C-terminal half is highly disordered^65^. RAD51 and RPA binding regions are also annotated^55,57^. **(c)** Net charge per residue as a function of residue index for RAD52 as calculated by the CIDER software from R. Pappu’s lab^66^. **(d)** Bright-field images of RAD52 mixtures under different protein and crowding conditions. **(e)** An example of Farid filter application to detect droplet edges in a bright field image, which is used to quantify the degree of droplet formation in images (see methods). **(f)** Quantitative phase separation state diagram of RAD52-PEG mixtures (see **Fig. S1**). **(g)** RAD52 droplet formation as a function of sodium chloride. **(h)** RAD52 droplet formation as a function of Calcium Chloride. **(i)** RAD52 droplet formation as a function of ATP. In **(g-i)**, the dashed line represents a non-phase separating sample. Bright-field images are shown in Figure S2.

Previous studies have shown that the structured ring of RAD52 N-terminal region constitutes two binding sites for ssDNA^67^. In addition, the C-terminal region of RAD52 binds RAD51 and RPA^57,68^, which are two key players in homologous recombination and other DNA repair pathways. Therefore, we tested whether RAD52 condensates can recruit these biomolecules and concentrate them. We measured the partition coefficient (droplet intensity/dilute phase intensity) of fluorescently labeled 40 nt poly(T) ssDNA (referred to as dT40 thereafter), Telomeric repeat-containing RNA (TERRA)^69^, RPA, and RAD51 within RAD52 condensates. We found that RAD52 condensates recruited ssDNA, RNA, RAD51, and RPA(**Fig. 2a-d**) with the partition coefficients all significantly larger than 1 (**Fig. 2e**). This indicates that RAD52 condensates have the potential to constitute a DNA repair hub that forms at ssDNA overhangs and recruits RPA, RAD51, and RNA. Condensation of RAD52 may underlie the formation of RAD52 repair centers, which were previously observed in cancer cells upon DNA damage^21,29^.

**Figure 2.**
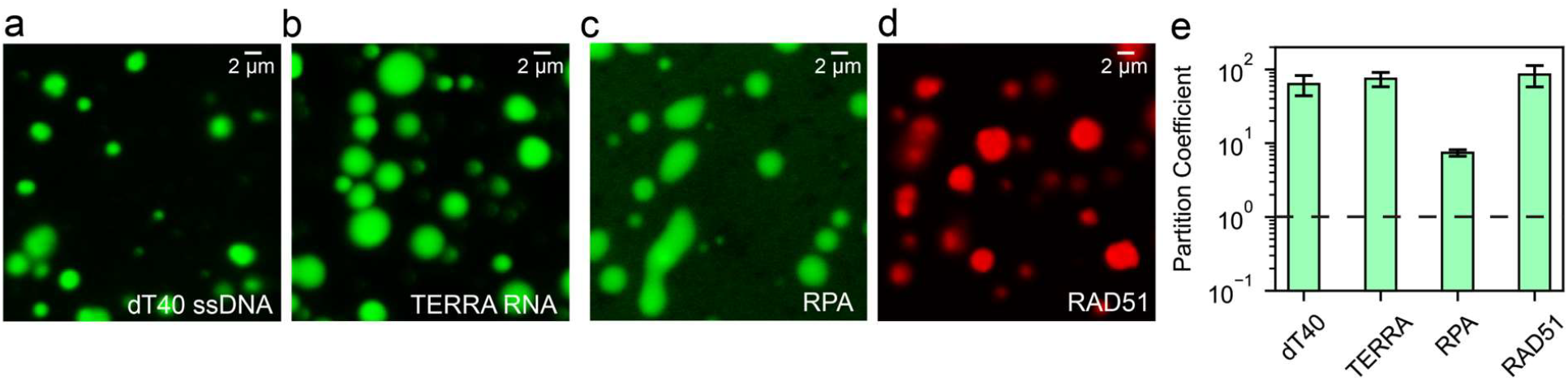
RAD52 condensates recruit DSB repair components. **(a-d)** Fluorescent images of RAD52 condensates (prepared at 5 µM RAD52) in the presence of dT40-Cy3, TERRA RNA-Cy3, RPA-MB543, and RAD51-Cy5, respectively. **(e)** Measured average partition coefficient of the client molecules (a-d) obtained from image analysis of multiple droplets per sample (n_condensates_>100). Error bars represent one standard deviation. The dashed line represents a partition coefficient of 1, indicating no preferential interactions between the client molecule and the droplet.

### ssDNA and RPA regulate RAD52 condensation

In a cellular context, changes in expression levels and accompanying changes in protein stoichiometries allow for concentration-based regulation of protein self-assembly^47,70^. Previous studies showed that heterotypic phase separation is generally governed by the stoichiometry of the interacting components^59,71,72^. Since RAD52 has binding domains for ssDNA and other repair proteins, we hypothesized that RAD52 phase separation may be modulated by the presence of ssDNA and DSB repair components in a concentration-dependent manner. To test this idea, we prepared RAD52 at 2.5 µM concentration in the absence of polymer crowders and varied the concentration of ssDNA dT40 (**Fig. 3a&b and Fig. S3**). Of note, dT40 length matches the circumference of the RAD52 ring^55^. We observed that DNA enhanced condensate formation of RAD52 at low molar fraction ([dT40]/[RAD52]∼0.1) (**Fig. 3a&b and Fig. S3**). This ratio corresponds to one ssDNA molecule per RAD52 undecameric ring^55^. However, at higher ssDNA-to-RAD52 ratios (>0.5) phase separation is completely abrogated (**Fig. 3a&b and Fig. S3**). This non-monotonic behavior, referred to as reentrant phase transition because the molecule re-enters the homogenous solution phase^59,73^, has been observed previously in mixtures of cationic proteins with nucleic acids and is thought to be a general feature of heterotypic phase separation mixtures^59,71,74,75^. We next tested the effect of RPA on RAD52 phase separation, since RPA is an interaction partner with RAD52. We found that RPA increased the tendency of phase separation in RAD52 mixtures at low RPA-to-RAD52 mixing ratios (**Fig. 3c&d**). Unlike ssDNA, we did not observe a subsequent decrease in phase separation at high RPA-to-RAD52 mixing ratios. This suggests that the decondensation threshold (the mixing ratio at which condensates re-enter the mixed phase) may exist beyond our concentration range.

**Figure 3.**
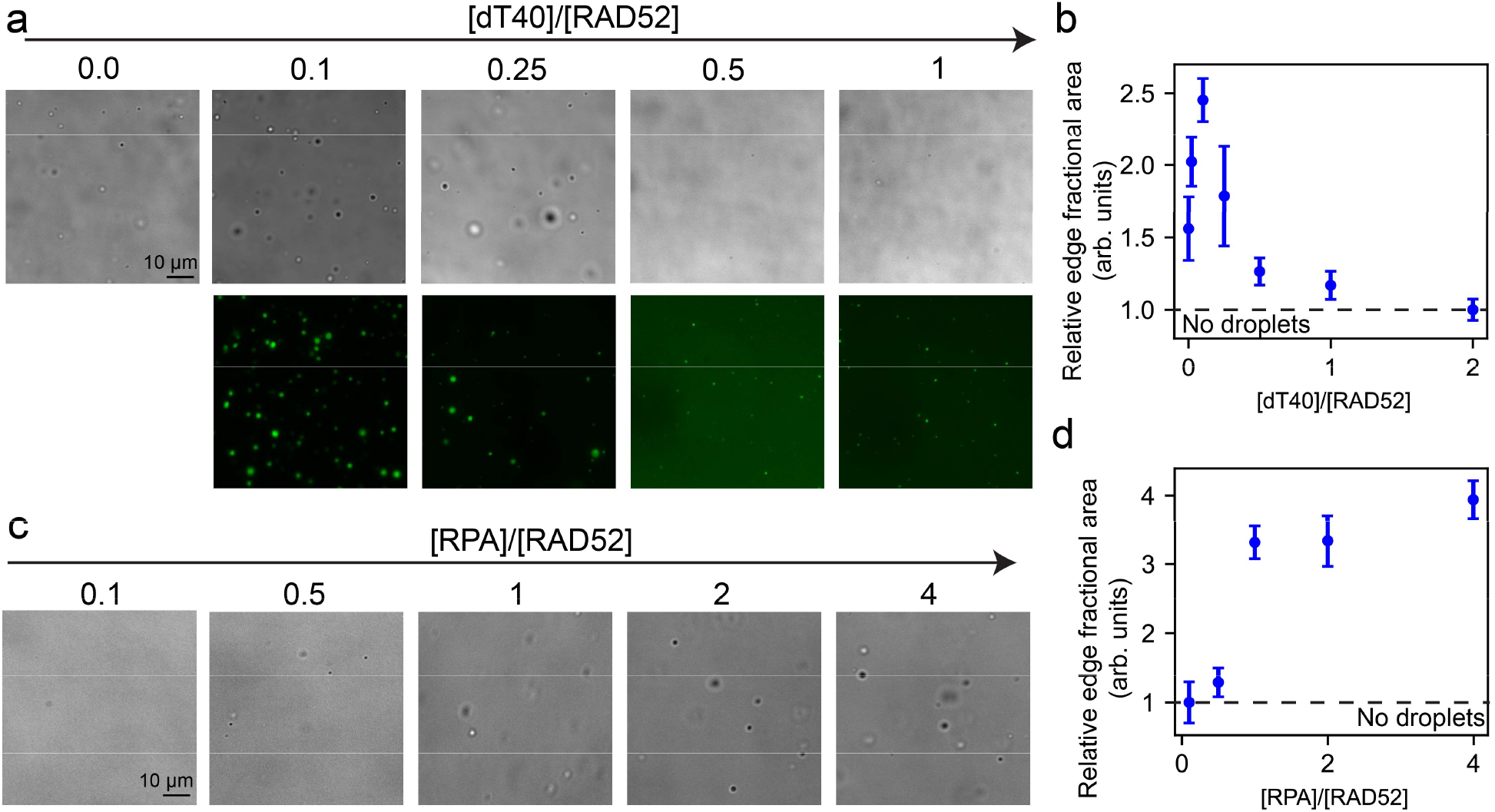
Regulation of RAD52 phase separation by ssDNA and RPA. **(a)** Bright-field and fluorescence images of RAD52-ssDNA mixtures at 2.5 µM RAD52 and variable dT40-to-RAD52 concentration ratio as indicated. dT40 in these experiments is labeled with Cy3 for fluorescence visualization. **(b)** Quantification of droplet formation via image analysis of the data in **(a)** and in **Fig. S3**. Dashed line indicates the no-droplet condition at 1. **(c)** Bright-field images of RAD52-RPA mixtures prepared at 2.5 µM RAD52 and variable RPA concentration. **(d)** Quantification of the images in **(c)** via measuring the interface fractional area normalized to a non-phase separating sample. The dashed line indicates the no-droplet condition at 1.

We next asked whether RAD52-ssDNA condensates and RAD52-RPA condensate form compatible phases with each other. Previous studies showed that multicomponent mixtures where multiple phases can form simultaneously may exist as one large mixed condensed phase or several distinct phases that are demixed from each other^76^. The formation of co-existing phases with distinct compositions is generally dictated by the hierarchy of interaction networks^77^. To test the compatibility of RAD52-ssDNA condensates and RAD52-RPA condensates, we formed each type separately (stained with 18-mer Cy5 conjugated ssDNA and RPA-MB543, see methods) and mixed them prior to imaging. Two-color imaging revealed that most of the condensates showed a mixture of RPA and RAD52 (**Fig. S4**). Accordingly, we conclude that a tri-component condensate comprised of RAD52, RPA, and ssDNA can be formed. Taken together, our experiment reveals that RAD52 phase separation is strongly modulated by RPA and ssDNA in a concentration-dependent manner. Importantly, RAD52 condensation is non-monotonically regulated by ssDNA through a reentrant phase transition. Such non-monotonic regulation may be an important physical property for the control of RAD52 condensation in the cell^78^. Conditions that alter intracellular concentrations such as RAD52 overexpression, increasing amounts of ssDNA (via end-resection), or elevated levels of RPA (upon significant DNA damage) may trigger RAD52 phase separation in the cell. Indeed, the formation of RAD52 foci has been observed in cells lacking BRCA2/BRCA1 proteins upon DNA damage^45^. We speculate that altered intracellular stoichiometry may modulate RAD52 condensation in the cell^16^.

### RAD52 induces RAD51-ssDNA fibril formation

Homologous recombination depends on the assembly of the RAD51 nucleoprotein complex on resected ssDNA overhangs to initiate the search for homology. The assembly of the RAD51 NPF depends on BRCA2 as it facilitates the replacement of RPA with RAD51 on ssDNA^17,79^. In BRCA2 mutated cells, the assembly of the NPF is thought to occur in a different pathway. To test if RAD52 phase separation affects the interactions between RAD51 and ssDNA, we prepared mixtures containing 5 µM RAD51 (with 10% RAD51-AF488) and 2.5 µM ssDNA (dT30) in a buffer devoid of ATP and divalent ions and did not find detectable structures (**Fig. 4a**). Adding RAD52 (5 µM) to the same mixture led to the formation of large fiber-like RAD51 structures that were more than 10 µm in length (**Fig. 4b**). Adding a crowding agent (5% PEG8K) resulted in additional formation of condensates that recruit RAD51 and form on RAD51 fibers (**Fig. 4b**). RAD51 fiber formation was observed even when RPA was present at a concentration equal to dT30 (2.5 µM, **Fig. 4c**), which is known to block RAD51-ssDNA interactions^60^. We then conducted three-color imaging experiments using fluorescently labeled RAD51, RAD52, and ssDNA. We observed that the fibrillar structures mainly contain RAD51 and ssDNA with a lower portion of RAD52, which is predominantly concentrated in spherical condensate-like structures from which RAD51 fibers appear to emanate (**Fig. 4d&e**). When the order of addition in the mixture is altered such that RAD52 is added before RAD51, we did not observe these fibrils (**Fig. S5**), suggesting that the fibrils form through interactions between RAD52 and the RAD51-ssDNA complex. Additionally, we confirmed that fibrillar structures can form even in the presence of ATP and calcium, which are necessary for the active form of the RAD51 recombinase (**Fig. S6**). Therefore, we hypothesized that RAD51 fibrils are formed via RAD52-induced clustering or bundling of RAD51 nucleoprotein filaments.

**Figure 4.**
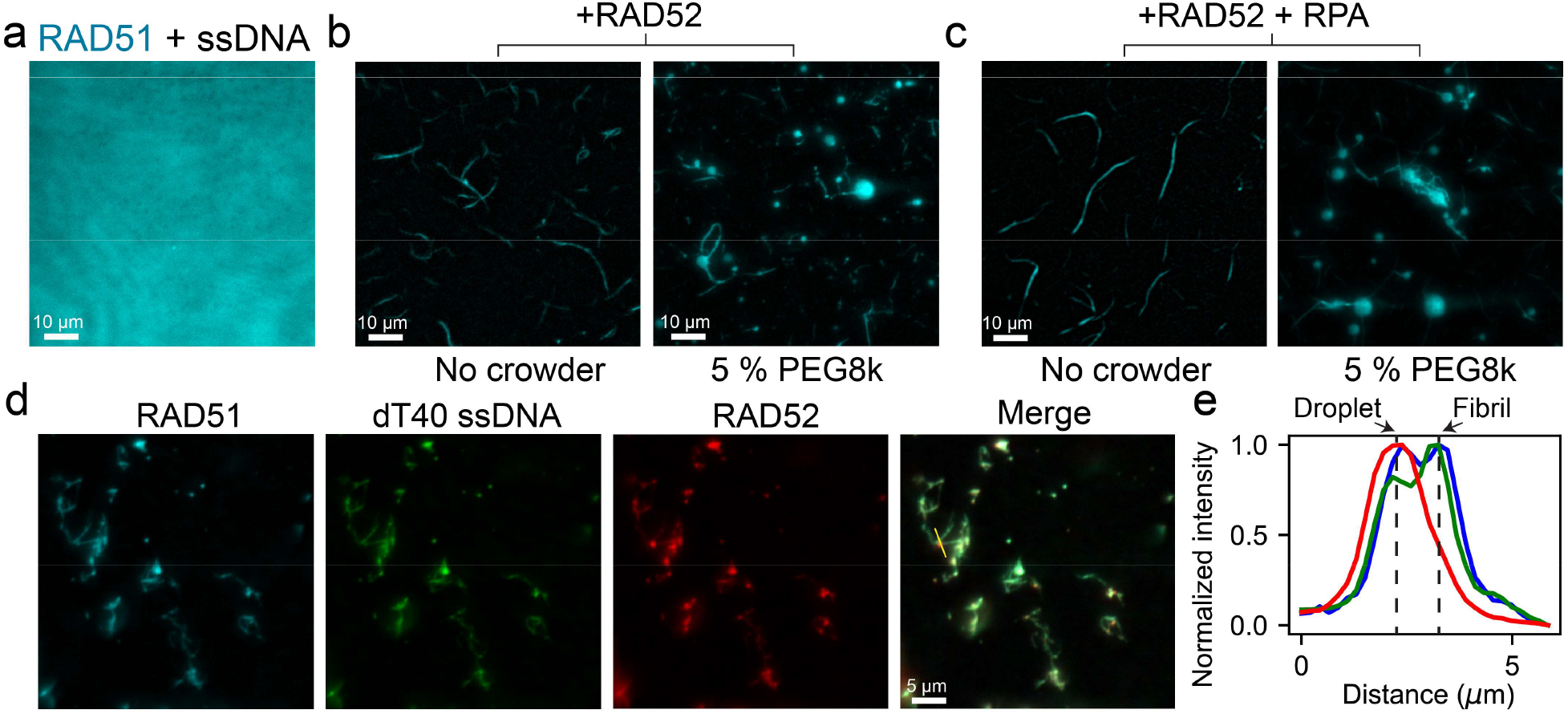
RAD52 induces RAD51-ssDNA fibril formation. **(a)** Fluorescent image of RAD51-ssDNA solution prepared at 5 µM RAD51 and 2.5 µM ssDNA dT30. Also added to the sample is ∼ 200 nM of AF488-conjugated RAD51. **(b)** Fluorescent images of the same sample in **(a)** but with an additional 5 µM of RAD52 in the presence and absence of a polymer crowder (PEG8k). **(c)** Fluorescent images of the same sample in **(b)** but with the addition of 2.5 µM RPA. **(d)** Three color fluorescent images of RAD51-RAD52-dT40 sample (5 µM, 5 µM, 2.5 µM, respectively) at no crowder condition. The sample contained ∼200 nM of Cy5-conjugated RAD52 and AF488-conjugated RAD51. **(e)** Intensity profile for the fluorescence signal across the yellow line in **(d)** for RAD51 (blue), dT40 (green), and RAD52 (red).

### Visualization of individual RAD51 nucleoprotein filaments using DNA-PAINT

Visualization of single RAD51 NPFs and their interactions with other proteins may provide significant insights into the mechanisms of RAD51 fiber formation observed herein. Imaging approaches to visualize the RAD51 NPF with nanometer resolution have only been achieved using Atomic Force Microscopy (AFM) and electron microscopy (EM)^80,81^. Both methods reveal limited information as they are label-free and cannot easily distinguish between different protein components. Single-molecule FRET experiments have been used to show that RAD51 filament formation extends the length of the DNA substrate^55^. Further, confocal microscopy has been used to visualize RAD51 filaments on dsDNA, albeit with diffraction-limited resolution^82^. The structure of RAD51 NPF has been previously visualized using cryo-electron microscopy^83^. However, aggregated or clustered filaments present insurmountable difficulties in cryo-EM imaging due to population heterogeneity and the large size of the supramolecular complexes. Furthermore, functional testing of the RAD51 NPF ability to invade homologous dsDNA templates has only been probed using gel-based assays and/or EM imaging^7,17^. In this section, we sought to develop an imaging approach that allows structural investigation of the RAD51 NPF in conjunctions with other proteins such as RAD52 and RPA as well as visualization of the D-loop formation with protein fluorescence staining.

To visualize RAD51 NPFs, D-loops, and NPF clustering, we applied a DNA-PAINT^84^ imaging approach with ∼10 nm resolution to a large DNA substrate, *ϕ*X174 virion DNA. The *ϕ*X174 bacteriophage genome is a 5.8 kb long ssDNA that is predominantly in circular form (∼80%) and is commercially available along with its dsDNA form. To assemble RAD51 NPFs, we mixed RAD51 with the ssDNA substrate at appropriate monovalent salt, divalent salt, and ATP concentrations (see methods) and kept the mixture at 37 °C for 1 hr (**Fig. 5a**). The assembled NPFs were then deposited on a poly(lysine)-coated microscope coverslip such that the complexes adhere to the coverslip surface. Protein visualization was done via immunostaining using primary antibodies against RAD51 and fragment secondary antibodies conjugated to probe oligonucleotides with the appropriate sequences for PAINT imaging (see methods). After immunostaining, a solution of fluorescently conjugated imager strands that are complementary to the probe strands on the secondary antibodies is used for imaging with appropriate concentrations to ensure rapid fluorescence blinking for subsequent image reconstruction. Placing the sample on an objective-TIRF microscope, we carried out DNA-PAINT imaging. Our approach yields imaging of single NPF complexes that are resolvable with a resolution of ∼15 nm (**Fig. 5b** and **Fig S7**). We could observe multiple circular RAD51 filaments in one field of view as well as several instances of linear RAD51 filaments (**Fig. 5b**). We then asked whether our super-resolution approach could detect D-loop formation. We mixed NPFs pre-assembled on *ϕ*X174 ssDNA with a homologous dsDNA target (*ϕ*X174 RF I) and used RPA to stain the ssDNA portion of the D-loop (**Fig. 5c**). If D-loop occurs, we expect to see significant colocalization between RPA and RAD51, however, due to the increased flexibility of RPA-ssDNA, we do not expect to resolve the chain contour using our approach. When preparing the NPFs without the dsDNA target, we observed that most NPFs were devoid of any RPA (**Fig. 5d**). However, when the NPFs were incubated with dsDNA, we observed significant co-localization between the RAD51 NPFs and RPA (**Fig. 5e**), indicating the formation of D-loops (**Fig. 5c**).

**Figure 5.**
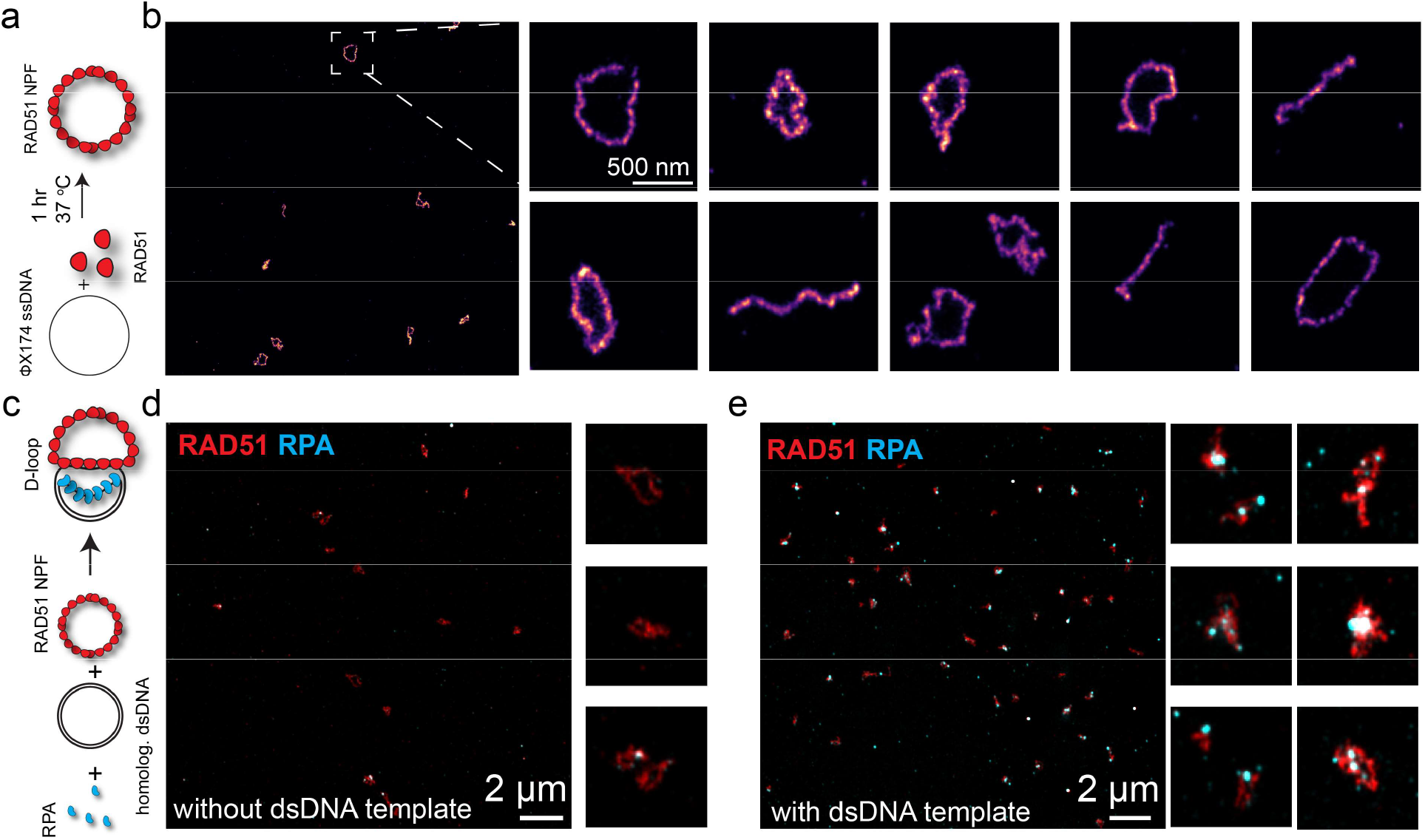
Super-resolution imaging to visualize RAD51 nucleoprotein filaments. **(a)** A scheme showing the assembly reaction of RAD51 NPF. **(b)** A representative large view of RAD51 NPFs on a coverslip surface reconstructed from DNA-PAINT imaging. Also shown are zoomed-in views of individual RAD51 NPFs. **(c)** A scheme showing the D-loop formation reaction in the presence of a homologous dsDNA template. **(d)** Two-color DNA-PAINT image of RAD51 NPFs (red) and RPA (cyan) in the absence of dsDNA template. Also shown are zoomed-in views of individual RAD51 NPFs. **(e)** Two-color DNA-PAINT image of RAD51 (red) and RPA (cyan) in the presence of a dsDNA template. The colocalization of the two proteins indicates the formation of D-loops. Also shown are zoomed-in views of individual D-loop structures.

### RAD52 induces clustering and end-to-end tethering of RAD51 nucleoprotein filaments

To probe the interactions between RAD52 and RAD51 NPFs, we next carried out the RAD51 NPF assembly reaction in the presence and absence of RAD52 and conducted super-resolution imaging with DNA-PAINT. All samples contained 1 µM RPA to ensure structure-free DNA templates. In the absence of RAD52, RAD51 NPFs were spread evenly on the coverslip surface and consistently existed as a single filament without clustering (**Fig. 6a, left**). In contrast, when RAD52 was added to the reaction mixture, we observed clusters of RAD51 NPFs (**Fig. 6a, right**). This clustering of NPFs is likely due to the RAD52 complexes acting as a hub to attract individual NPFs via their multiple IDRs. We also observed long NPFs that emanate from the clusters (**Fig. 6a, right**). These long filaments constituted ∼40% of all detected filaments (**Fig 6b**). The overall length of RAD51 NPFs is much larger in the presence of RAD52 (9 ± 6 µm) than in its absence (2.1 ± 0.4 µm, **Fig. 6c**). Importantly, these filaments were observed at multiple RPA concentrations (**Fig. S8**). The presence of long RAD51 NPF was also observed when the NPFs were assembled on the dsDNA substrate, although with overall shorter filaments (**Fig. S9**). We suspected that these exceptionally long filaments are individual linear filaments tethered to each other via RAD52. To test this, we performed two-color DNA PAINT imaging to inspect whether RAD52 exists in these clusters. Indeed, our two-color imaging reveals that RAD52 was present in RAD51 NPF clusters, especially at the junction points (**Fig. 6d**). We also observed instances where multiple curved filaments are attached with points of attachments containing localizations of RAD52 (**Fig. 6e**). Importantly, we observed RAD52 localizations that are evenly spaced on the linear elongated RAD51 NPFs (**Fig. 6d&f** and **Fig. S10**). The distance between successive RAD52 spots agrees with the measured length of individual RAD51 filaments formed on the ssDNA substrate (**Fig. 6f & Fig. S10**). These results indicate that RAD52 has two modes of interaction with RAD51 NPFs that facilitate fiber formation. The first is RAD52 bridging multiple filaments together side by side (**Fig. 6d&e**). The second mode is RAD52 facilitating end-to-end tethering of RAD51 NPFs (**Fig. 6d&f** and **Fig. S10**). A combination of these two modes of interactions can explain the RAD51-ssDNA fibril formation observed in this work. Note that we do not see large DNA bundles in these experiments because of the need to use extremely low concentrations of components that allow us to resolve DNA structures as well as the use of larger DNA substrates that are slower in diffusion. Taken together, our super-resolution imaging experiments reveal that RAD52 can induce clustering and end-to-end tethering of RAD51 NPFs.

**Figure 6.**
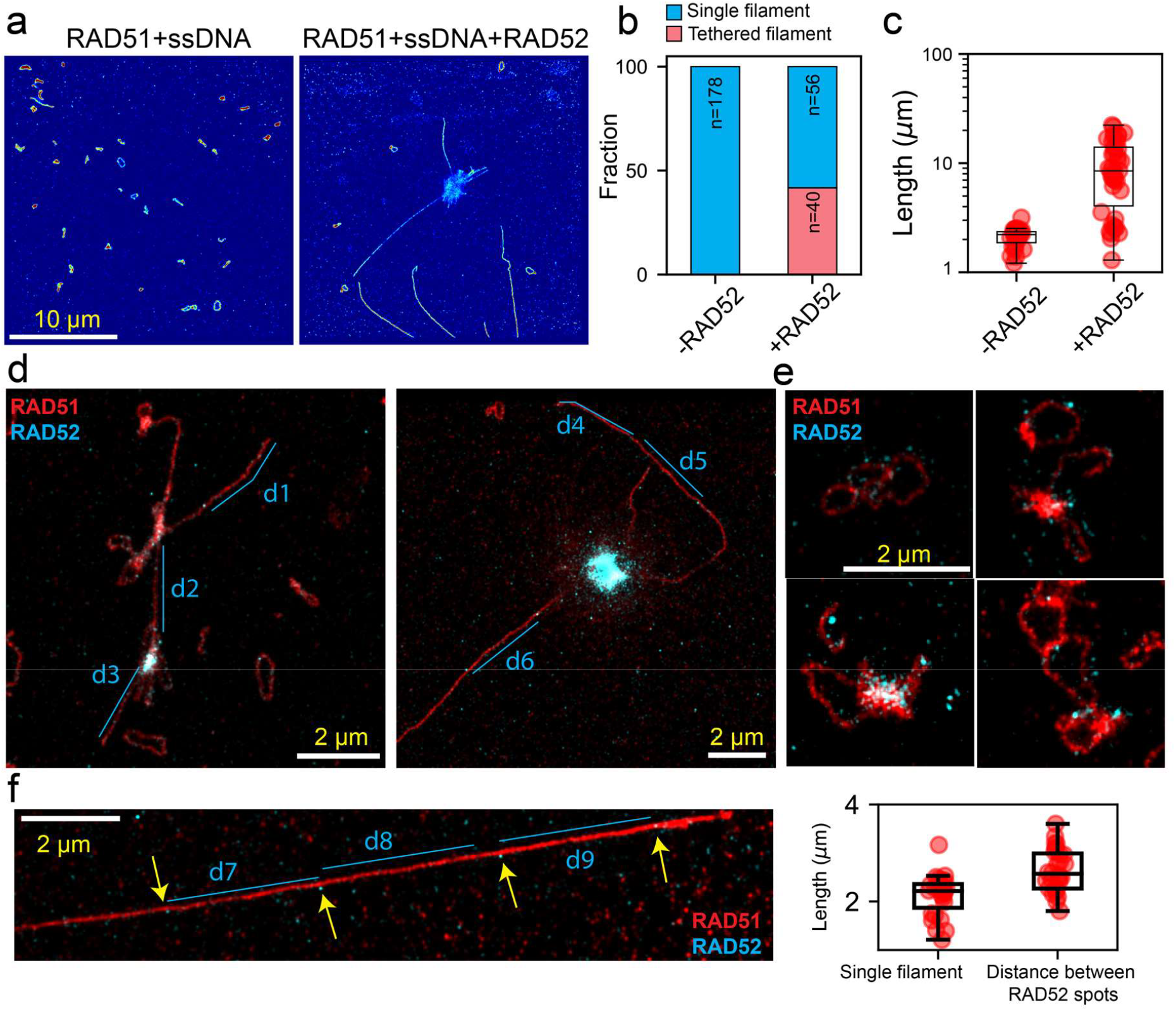
RAD52 induces RAD51 nucleoprotein filament clustering and end-to-end tethering. **(a)** DNA-PAINT super-resolution image of RAD51- nucleoprotein filaments formed on *ϕ*x174 ssDNA virion in the absence of RAD52 (left) and the presence of 0.5 µM RAD52. **(b)** Quantification of the single-filaments and the tethered filaments in panel **(a)** obtained from analyzing multiple images. **(c)** DNA-PAINT image of RAD52-induced clustering of RAD51 NPF assembled on dsDNA. **(d)** Two representative DNA-PAINT images of RAD52 (cyan) and RAD51 NPF (red) clusters showing elongated RAD51 NPFs that are likely to be tethered (also see **Fig. S10**). **(e)** Two-color images of clustered circular RAD51 NPFs (red) in the presence of RAD52 (cyan). **(f)** Example of an excessively long RAD51 NPF with equally spaced localizations of RAD52 (cyan, yellow arrows). The box plot on the right shows the measured distance between RAD52 spots in the left image, **(d)**, and **Fig. S10**. The RAD52 inter-spot distance is compared with the measured length of single RAD51 NPFs from the data in **(a)**.

To further test our findings from super-resolution imaging, we utilized atomic force microscopy (AFM) to image RAD51 NPFs and RAD52-RAD51 complexes without the use of fluorophore labeling or antibody staining. First, we imaged the single-stranded DNA template, confirming the presence of both circular and linear chains (**Fig. 7a**). Next, we examined pre-assembled RAD51 filaments and confirmed that these filaments existed separately without inter-filament clustering (**Fig 7b**). When RAD51 NPFs were assembled in the presence of RAD52, we observed large-scale filaments (∼3-5 µm) in addition to single filaments that were longer than those observed in the absence of RAD52 (∼2 µm, **Fig. 7c&d**). This is evidenced by a wider distribution of filament lengths (**Fig. 7d**). These long filaments were interspersed with large blobs that we identify as RAD52, since RAD52 rings are ∼10 nm in diameter and are much larger than RAD51. Notably, in some cases, we observed filaments that were too long to be captured within one field of view (4×4 µm for our AFM setup). We also found networks of multiple long RAD51 filaments that are apparently crosslinked by RAD52 blobs (**Fig. 7c, bottom**). In sum, our AFM imaging results corroborate our DNA-PAINT findings and point to a unique ability of RAD52 to induce clustering and end-to-end tethering of RAD51 NPFs (**Fig. 7e**).

**Figure 7.**
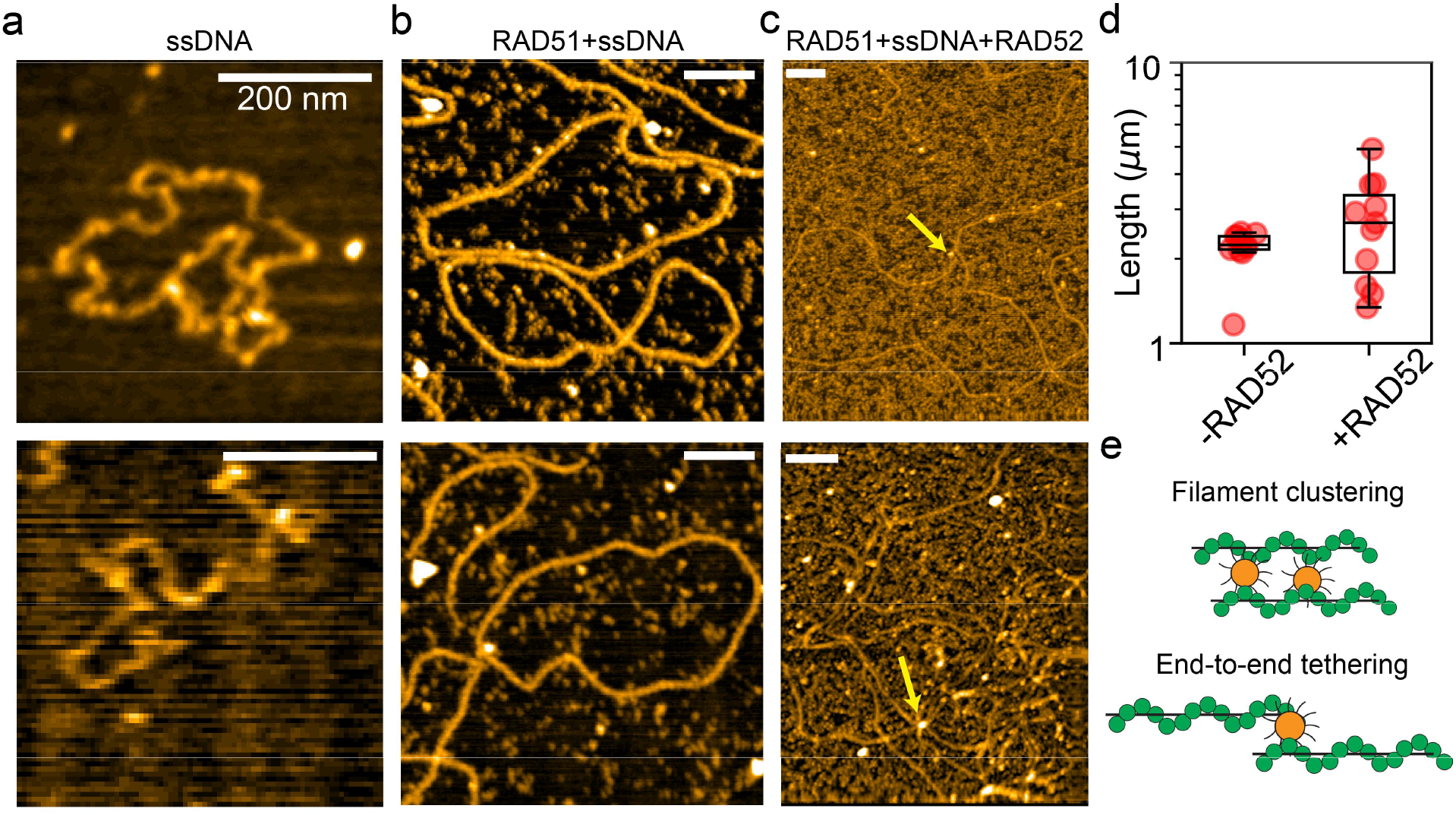
Atomic Force microscopy confirms RAD52-induced clustering and tethering of RAD51 NPFs. **(a)** AFM images of naked *ϕ*X174 ssDNA in its circular (top) and linear (bottom) forms. **(b)** AFM images of RAD51 NPF. **(c)** AFM image of RAD51 NPFs in the presence of 0.5 µM of RAD52. All scale bars are 200 nm. Arrows indicate a molecule that is likely to be RAD52 crosslinking filaments. **(d)** Quantification of filament length from AFM images showing extended lengths for RAD51 NPFs in the presence of RAD52. Larger filament lengths were not obtainable due to the small field of view in AFM (4×4 µm). **(e)** Scheme showing the two bridging modes of RAD52 that can either cluster or tether RAD51 filaments.

## Discussion and Conclusion

The human RAD52 complex holds great promise for cancer therapy as its disruption is exclusively toxic to cells that lack functional BRCA2 gene^70^, which underlies many types of cancer as well as development disorders that are collectively called homologous recombination deficiency syndrome^85^. A significant amount of research has been devoted to showing how this protein complex interacts with ssDNA and cataloging its biochemical activities such as single-stranded DNA annealing and reverse strand exchange reactions^35,44^. However, the self-assembly properties of the human RAD52 and its interaction with RAD51 NPFs have not been fully documented. Here, we filled this gap by studying the self-assembly properties of RAD52 and its interactions with the RAD51 nucleoprotein filament, which is the central macromolecular machine in the HR pathway. We show that RAD52 undergoes phase separation at sub-micromolar concentrations in crowded environments. These RAD52 condensates recruit nucleic acids and DSB repair proteins, potentiating them to serve as a model for the RAD52 repair centers that have been reported in the literature. We further show that RAD52 induces the formation of RAD51-ssDNA fibril like structures that span tens of microns in length. Simultaneously, single molecule imaging shows that RAD52 induces clustering of RAD51 nucleoprotein filaments as well as end-to-end tethering of linear RAD51 nucleoprotein filaments. Taken together, our findings show unique organizational features of the RAD52 complex that are likely encoded by its branched multivalent structure.

The phase separation properties of RAD52 may underlie RAD52 foci formation in cells that act as DSB repair centers by recruiting RAD51 and ssDNA^86^. This adds to the emerging paradigm of linking phase separation to DNA damage processes in the cell^87^. A recent report showed that yeast Rad52 can phase separate in the nucleus and induce the formation of nuclear microtubules^88^. It was also reported recently that RAD51 itself can form amyloid fibrils at high concentrations^89^. Importantly, two studies in the literature have found ∼ 1 µm thick RAD51 fibril-like structures in human cells upon RAD51 overexpression^89,90^. These structures^90^ show a stark similarity with the RAD52-induced fibrils shown in **Figure 4**. Another study showed that RAD51 binding domain TR2 of BRCA2 induces bundling of RAD51-ssDNA NPFs in vitro^79^. This specific domain is required for the BRCA2 function in protecting stalled replication forks^91^. Our study shows a similar effect of RAD52-induced bundling of RAD51-ssDNA complexes. Interestingly, RAD52 is also known to be important in protecting stalled replication forks^37^. Another important work showed that RecA protein, which is the RAD51 equivalent in *E*.*coli*, forms RecA bundles/fibers upon DNA damage^92^. Remarkably, the bundling of RecA filaments includes ssDNA-recA filaments as well as DNA-free recA filaments. It was therefore proposed that RecA bundles provide a mechanism to channel the movement of ssDNA along the RecA fiber through filament sliding, which allows pairing between the ssDNA overhang and the homology locus. The human RAD51 fibers we observed appear similar to those observed for the RecA protein^92^ and upon RAD51 overexpression in human cells^90^. Further effort must be dispensed into understanding the physiological relevance of RAD51 fibrils and whether they contribute to later stages of HR such as homology search.

Our single molecule imaging showed a unique ability of RAD52 to tether pre-formed RAD51 NPFs from their ends, leading to the formation of exceptionally long presynaptic filaments. This observation is consistent with a previous report showing that the human RAD52 complex has an intrinsic preference for binding DNA ends^93^, similar to the Ku heterodimer in the NHEJ pathway^94^. We speculate this feature of the RAD52 complex contributes to its ability to promote second-end capture^95,96^. When a DSB occurs, two RAD51 filaments form on the two ends of the DSB. After strand invasion, the ssDNA portion of the D-loop can capture the other RAD51 filament to form a joint protein-DNA complex where the two RAD51 filaments bind to both complementary strands of the same dsDNA donor. This phenomenon is known as second-end capture and is promoted by RAD52^95,96^. We argue that the ability of RAD52 to tether RAD51 NPF ends might contribute to its capacity to promote second-end capture by ensuring both DSB-induced RAD51 NPF remain together physically during homology search.

In conclusion, we have investigated the self-assembly properties of the human RAD52 complex and its interactions with DSB repair proteins and nucleic acids. We show that RAD52 can undergo phase separation, induce RAD51-ssDNA fibril formation, and facilitate RAD51 NPF clustering and end-to-end tethering. These properties may provide a mechanistic basis for the protein function in the cell, which may ultimately lead to advancing our current strategies of inhibiting RAD52 that are harnessed in drug-based cancer therapy.

## Author Contributions

I.A. conceived the idea and designed the study with input from T.H. & M.S. I.A. curated and analyzed the data with help from P.A.D. S.P. and T.W.L conducted the AFM imaging and analysis. M.H. and M.S. expressed and purified the proteins. I.A. wrote the manuscript with input from S.P., T.H., and M.S. All authors revised the manuscript.

## Acknowledgments

The authors are grateful to Edwin Anthony’s lab for graciously providing site-specifically labeled RPA protein and for useful discussions. T. Ha is a Howard Hughes Medical Institute (HHMI) investigator. T. Ha acknowledges funding from the National Institutes of Health (NIH/NIGMS R35 GM122569). M. Spies acknowledges support from the National Cancer Institute (NIH/NCI R01CA232425). I. Alshareedah acknowledges support from HHMI and the Jane Coffin Childs Memorial Fund for Medical Research.

## Competing Interests

The authors declare no competing interests.

## Material and Data Availability

All the data pertaining to the conclusions of this manuscript are presented in the main text and supplementary information. Additional data are available upon reasonable request.

## Notes

### Competing Interest Statement

The authors have declared no competing interest.

## References

1 Tubbs, A. & Nussenzweig, A. Endogenous DNA damage as a source of genomic instability in cancer. Cell 168, 644–656 (2017).

2 McKinnon, P. J. & Caldecott, K. W. DNA strand break repair and human genetic disease. Annu. Rev. Genomics Hum. Genet. 8, 37–55 (2007).

3 Woodward, E. R. & Meyer, S. Fanconi Anaemia, Childhood Cancer and the BRCA Genes. Genes 12, 1520 (2021).

4 Cannan, W. J. & Pederson, D. S. Mechanisms and consequences of double-strand DNA break formation in chromatin. Journal of cellular physiology 231, 3–14 (2016).

5 Mao, Z., Bozzella, M., Seluanov, A. & Gorbunova, V. DNA repair by nonhomologous end joining and homologous recombination during cell cycle in human cells. Cell cycle 7, 2902–2906 (2008).

6 Krejci, L., Altmannova, V., Spirek, M. & Zhao, X. Homologous recombination and its regulation. Nucleic acids research 40, 5795–5818 (2012).

7 Zhao, X. et al. Cell cycle-dependent control of homologous recombination. Acta biochimica et biophysica Sinica 49, 655–668 (2017).

8 Kimble, M. T., Johnson, M. J., Nester, M. R. & Symington, L. S. Long-range DNA end resection supports homologous recombination by checkpoint activation rather than extensive homology generation. Elife 12, e84322 (2023).

9 Wang, Y.-L. et al. MRNIP condensates promote DNA double-strand break sensing and end resection. Nature communications 13, 2638 (2022).

10 Cejka, P. & Symington, L. S. DNA End Resection: Mechanism and Control. Annual Review of Genetics 55, 285–307 (2021). 10.1146/annurev-genet-071719-020312

11 Li, X. & Heyer, W.-D. Homologous recombination in DNA repair and DNA damage tolerance. Cell research 18, 99–113 (2008).

12 Wold, M. S. Replication protein A: a heterotrimeric, single-stranded DNA-binding protein required for eukaryotic DNA metabolism. Annual review of biochemistry 66, 61–92 (1997).

13 Sung, P. & Klein, H. Mechanism of homologous recombination: mediators and helicases take on regulatory functions. Nature reviews Molecular cell biology 7, 739–750 (2006).

14 Haber, J. E. DNA repair: the search for homology. Bioessays 40, 1700229 (2018).

15 Bhat, K. P. & Cortez, D. RPA and RAD51: fork reversal, fork protection, and genome stability. Nature structural & molecular biology 25, 446–453 (2018).

16 Andriuskevicius, T., Kotenko, O. & Makovets, S. Putting together and taking apart: assembly and disassembly of the Rad51 nucleoprotein filament in DNA repair and genome stability. Cell Stress 2, 96 (2018).

17 Jensen, R. B., Carreira, A. & Kowalczykowski, S. C. Purified human BRCA2 stimulates RAD51-mediated recombination. Nature 467, 678–683 (2010).

18 Andreassen, P. R., Seo, J., Wiek, C. & Hanenberg, H. Understanding BRCA2 Function as a Tumor Suppressor Based on Domain-Specific Activities in DNA Damage Responses. Genes 12, 1034 (2021).

19 Sung, P. Function of yeast Rad52 protein as a mediator between replication protein A and the Rad51 recombinase. Journal of Biological Chemistry 272, 28194–28197 (1997).

20 Nogueira, A., Fernandes, M., Catarino, R. & Medeiros, R. RAD52 functions in homologous recombination and its importance on genomic integrity maintenance and cancer therapy. Cancers 11, 1622 (2019).

21 Feng, Z. et al. Rad52 inactivation is synthetically lethal with BRCA2 deficiency. Proceedings of the National Academy of Sciences 108, 686–691 (2011).

22 McIlwraith, M. J. et al. Reconstitution of the strand invasion step of double-strand break repair using human Rad51 Rad52 and RPA proteins. Journal of molecular biology 304, 151–164 (2000).

23 Benson, F. E., Baumann, P. & West, S. C. Synergistic actions of Rad51 and Rad52 in recombination and DNA repair. Nature 391, 401–404 (1998).

24 Muhammad, A. A. et al. Human RAD52 stimulates the RAD51-mediated homology search. Life Science Alliance 7 (2024).

25 Hanamshet, K., Mazina, O. M. & Mazin, A. V. Reappearance from obscurity: mammalian Rad52 in homologous recombination. Genes 7, 63 (2016).

26 Bhat, D. S., Spies, M. A. & Spies, M. A moving target for drug discovery: Structure activity relationship and many genome (de)stabilizing functions of the RAD52 protein. DNA Repair 120, 103421 (2022). 10.1016/j.dnarep.2022.103421

27 Toma, M., Sullivan-Reed, K., Śliwiński, T. & Skorski, T. RAD52 as a potential target for synthetic lethality-based anticancer therapies. Cancers 11, 1561 (2019).

28 Mahajan, S., Raina, K., Verma, S. & Rao, B. Human RAD52 protein regulates homologous recombination and checkpoint function in BRCA2 deficient cells. The international journal of biochemistry & cell biology 107, 128–139 (2019).

29 Lok, B., Carley, A., Tchang, B. & Powell, S. RAD52 inactivation is synthetically lethal with deficiencies in BRCA1 and PALB2 in addition to BRCA2 through RAD51-mediated homologous recombination. Oncogene 32, 3552–3558 (2013).

30 Hanamshet, K. & Mazin, A. V. The function of RAD52 N-terminal domain is essential for viability of BRCA-deficient cells. Nucleic Acids Research 48, 12778–12791 (2020). 10.1093/nar/gkaa1145

31 Wang, Z. et al. Association of Germline BRCA2 Mutations With the Risk of Pediatric or Adolescent Non–Hodgkin Lymphoma. JAMA Oncology 5, 1362–1364 (2019). 10.1001/jamaoncol.2019.2203

32 Petrucelli, N., Daly, M. B. & Pal, T. BRCA1-and BRCA2-associated hereditary breast and ovarian cancer. (2022).

33 Lancaster, J. M. et al. BRCA2 mutations in primary breast and ovarian cancers. Nature genetics 13, 238–240 (1996).

34 Schuler, N. et al. DNA-Damage Foci to Detect and Characterize DNA Repair Alterations in Children Treated for Pediatric Malignancies. PLOS ONE 9, e91319 (2014). 10.1371/journal.pone.0091319

35 Reddy, G., Golub, E. I. & Radding, C. M. Human Rad52 protein promotes single-strand DNA annealing followed by branch migration. Mutation Research/Fundamental and Molecular Mechanisms of Mutagenesis 377, 53–59 (1997).

36 Rothenberg, E., Grimme, J. M., Spies, M. & Ha, T. Human Rad52-mediated homology search and annealing occurs by continuous interactions between overlapping nucleoprotein complexes. Proceedings of the National Academy of Sciences 105, 20274–20279 (2008).

37 Malacaria, E. et al. Rad52 prevents excessive replication fork reversal and protects from nascent strand degradation. Nature Communications 10, 1412 (2019). 10.1038/s41467-019-09196-9

38 Murfuni, I. et al. Survival of the Replication Checkpoint Deficient Cells Requires MUS81-RAD52 Function. PLOS Genetics 9, e1003910 (2013). 10.1371/journal.pgen.1003910

39 Sotiriou, S. K. et al. Mammalian RAD52 functions in break-induced replication repair of collapsed DNA replication forks. Molecular cell 64, 1127–1134 (2016).

40 Min, J., Wright, W. E. & Shay, J. W. Clustered telomeres in phase-separated nuclear condensates engage mitotic DNA synthesis through BLM and RAD52. Genes & development 33, 814–827 (2019).

41 Jalan, M., Olsen, K. S. & Powell, S. N. Emerging roles of RAD52 in genome maintenance. ancers 11, 1038 (2019).

42 Yasuhara, T. et al. Human Rad52 promotes XPG-mediated R-loop processing to initiate transcription-associated homologous recombination repair. Cell 175, 558–570.e511 (2018).

43 Liu, S. et al. DNA repair protein RAD52 is required for protecting G-quadruplexes in mammalian cells. Journal of Biological Chemistry 299 (2023).

44 Mazina, O. M., Keskin, H., Hanamshet, K., Storici, F. & Mazin, A. V. Rad52 inverse strand exchange drives RNA-templated DNA double-strand break repair. Molecular cell 67, 19–29.e13 (2017).

45 Alvaro, D., Lisby, M. & Rothstein, R. Genome-wide analysis of Rad52 foci reveals diverse mechanisms impacting recombination. PLoS genetics 3, e228 (2007).

46 Lok, B. H. & Powell, S. N. Molecular Pathways: Understanding the Role of Rad52 in Homologous Recombination for Therapeutic AdvancementThe Role of Rad52 in Homologous Recombination. Clinical cancer research 18, 6400–6406 (2012).

47 Ho, V. et al. Aberrant expression of RAD52, its prognostic impact in rectal cancer and association with poor survival of patients. International Journal of Molecular Sciences 21, 1768 (2020).

48 Liu, Y., Li, M.-j., Lee, E. Y. P. & Maizels, N. Localization and dynamic relocalization of mammalian Rad52 during the cell cycle and in response to DNA damage. Current biology 9, 975–978 (1999).

49 Kinoshita, C. et al. The cryo-EM structure of full-length RAD52 protein contains an undecameric ring. FEBS Open Bio 13, 408–418 (2023).

50 Saotome, M. et al. Structural Basis of Homology-Directed DNA Repair Mediated by RAD52. iScience 3, 50–62 (2018). 10.1016/j.isci.2018.04.005

51 Kagawa, W. et al. Crystal Structure of the Homologous-Pairing Domain from the Human Rad52 Recombinase in the Undecameric Form. Molecular Cell 10, 359–371 (2002). 10.1016/S1097-2765(02)00587-7

52 Singleton, M. R., Wentzell, L. M., Liu, Y., West, S. C. & Wigley, D. B. Structure of the single-strand annealing domain of human RAD52 protein. Proceedings of the National Academy of Sciences 99, 13492–13497 (2002). 10.1073/pnas.212449899

53 Liang, C.-C. et al. Mechanism of single-stranded DNA annealing by RAD52–RPA complex. Nature 629, 697–703 (2024). 10.1038/s41586-024-07347-7

54 Honda, M. et al. Human RAD52 double-ring remodels replication forks restricting fork reversal. bioRxiv, 2023.2011.2014.566657 (2023). 10.1101/2023.11.14.566657

55 Grimme, J. M. et al. Human Rad52 binds and wraps single-stranded DNA and mediates annealing via two hRad52–ssDNA complexes. Nucleic acids research 38, 2917–2930 (2010).

56 Khade, N. V. & Sugiyama, T. Roles of C-terminal region of yeast and human rad52 in rad51-nucleoprotein filament formation and ssDNA annealing. PLoS One 11, e0158436 (2016).

57 Shen, Z., Cloud, K. G., Chen, D. J. & Park, M. S. Specific Interactions between the Human RAD51 and RAD52 Proteins (*). Journal of Biological Chemistry 271, 148–152 (1996).

58 Wilken, S., Chaderjian, A. & Saleh, O. A. Spatial organization of phase-separated DNA droplets. Physical Review X 13, 031014 (2023).

59 Banerjee, P. R., Milin, A. N., Moosa, M. M., Onuchic, P. L. & Deniz, A. A. Reentrant phase transition drives dynamic substructure formation in ribonucleoprotein droplets. Angewandte Chemie 129, 11512–11517 (2017).

60 Ma, C. J., Gibb, B., Kwon, Y., Sung, P. & Greene, E. C. Protein dynamics of human RPA and RAD51 on ssDNA during assembly and disassembly of the RAD51 filament. Nucleic acids research 45, 749–761 (2017).

61 Handwerger, K. E., Cordero, J. A. & Gall, J. G. Cajal bodies, nucleoli, and speckles in the Xenopus oocyte nucleus have a low-density, sponge-like structure. Molecular biology of the cell 16, 202–211 (2005).

62 Farid, H. & Simoncelli, E. P. Diferentiation of discrete multidimensional signals. IEEE Transactions on image processing 13, 496–508 (2004).

63 Tomasello, G., Armenia, I. & Molla, G. The Protein Imager: a full-featured online molecular viewer interface with server-side HQ-rendering capabilities. Bioinformatics 36, 2909–2911 (2020). 10.1093/bioinformatics/btaa009

64 Balboni, B. et al. An integrative structural study of the human full-length RAD52 at 2.2 Å resolution. Communications Biology 7, 956 (2024). 10.1038/s42003-024-06644-1

65 Romero, P. et al. Sequence complexity of disordered protein. Proteins: Structure, Function, and Bioinformatics 42, 38–48 (2001).

66 Holehouse, A. S., Ahad, J., Das, R. K. & Pappu, R. V. CIDER: classification of intrinsically disordered ensemble regions. Biophysical journal 108, 228a (2015).

67 Kagawa, W. et al. Identification of a second DNA binding site in the human Rad52 protein. Journal of Biological Chemistry 283, 24264–24273 (2008).

68 Park, M. S., Ludwig, D. L., Stigger, E. & Lee, S.-H. Physical interaction between human RAD52 and RPA is required for homologous recombination in mammalian cells. Journal of Biological Chemistry 271, 18996–19000 (1996).

69 Luke, B. & Lingner, J. TERRA: telomeric repeat-containing RNA. The EMBO journal 28, 2503–2510 (2009).

70 Rossi, M. J., DiDomenico, S. F., Patel, M. & Mazin, A. V. RAD52: paradigm of synthetic lethality and new developments. Frontiers in Genetics 12, 780293 (2021).

71 Alshareedah, I. et al. Interplay between short-range attraction and long-range repulsion controls reentrant liquid condensation of ribonucleoprotein–RNA complexes. Journal of the American Chemical Society 141, 14593–14602 (2019).

72 Mittag, T. & Pappu, R. V. A conceptual framework for understanding phase separation and addressing open questions and challenges. Molecular cell 82, 2201–2214 (2022).

73 Alshareedah, I., Moosa, M. M., Raju, M., Potoyan, D. A. & Banerjee, P. R. Phase transition of RNA™ protein complexes into ordered hollow condensates. Proceedings of the National Academy of Sciences 117, 15650–15658 (2020).

74 Alshareedah, I., Thurston, G. M. & Banerjee, P. R. Quantifying viscosity and surface tension of multicomponent protein-nucleic acid condensates. Biophysical journal 120, 1161–1169 (2021).

75 Alshareedah, I., Moosa, M. M., Pham, M., Potoyan, D. A. & Banerjee, P. R. Programmable viscoelasticity in protein-RNA condensates with disordered sticker-spacer polypeptides. Nature communications 12, 6620 (2021).

76 Feric, M. et al. Coexisting liquid phases underlie nucleolar subcompartments. Cell 165, 1686–1697 (2016).

77 Jacobs, W. M. Theory and simulation of multiphase coexistence in biomolecular mixtures. Journal of Chemical Theory and Computation 19, 3429–3445 (2023).

78 Henninger, J. E. et al. RNA-mediated feedback control of transcriptional condensates. Cell 184, 207–225.e224 (2021).

79 Appleby, R., Joudeh, L., Cobbett, K. & Pellegrini, L. Structural basis for stabilisation of the RAD51 nucleoprotein filament by BRCA2. Nature Communications 14, 7003 (2023).

80 Ristic, D. et al. Human Rad51 filaments on double-and single-stranded DNA: correlating regular and irregular forms with recombination function. Nucleic acids research 33, 3292–3302 (2005).

81 Van Dyck, E., Hajibagheri, N. M., Stasiak, A. & West, S. C. Visualisation of human rad52 protein and its complexes with hRad51 and DNA. Journal of molecular biology 284, 1027–1038 (1998).

82 Gibb, B. et al. Protein dynamics during presynaptic-complex assembly on individual single-stranded DNA molecules. Nature structural & molecular biology 21, 893–900 (2014).

83 Short, J. M. et al. High-resolution structure of the presynaptic RAD51 filament on single-stranded DNA by electron cryo-microscopy. Nucleic acids research 44, 9017–9030 (2016).

84 Schnitzbauer, J., Strauss, M. T., Schlichthaerle, T., Schueder, F. & Jungmann, R. Super-resolution microscopy with DNA-PAINT. Nature protocols 12, 1198–1228 (2017).

85 Toh, M. & Ngeow, J. Homologous recombination deficiency: cancer predispositions and treatment implications. The Oncologist 26, e1526–e1537 (2021).

86 Essers, J. et al. Nuclear dynamics of RAD52 group homologous recombination proteins in response to DNA damage. The EMBO journal 21, 2030–2037 (2002).

87 Spegg, V. & Altmeyer, M. Biomolecular condensates at sites of DNA damage: More than just a phase. DNA repair 106, 103179 (2021).

88 Oshidari, R. et al. DNA repair by Rad52 liquid droplets. Nature communications 11, 695 (2020).

89 Kachkin, D. V. et al. Human RAD51 Protein Forms Amyloid-like Aggregates In Vitro. International Journal of Molecular Sciences 23, 11657 (2022).

90 Raderschall, E. et al. Formation of higher-order nuclear Rad51 structures is functionally linked to p21 expression and protection from DNA damage-induced apoptosis. Journal of cell science 115, 153–164 (2002).

91 Kim, T. M., Son, M. Y., Dodds, S., Hu, L. & Hasty, P. Deletion of BRCA2 exon 27 causes defects in response to both stalled and collapsed replication forks. Mutation Research/Fundamental and Molecular Mechanisms of Mutagenesis 766, 66–72 (2014).

92 Lesterlin, C., Ball, G., Schermelleh, L. & Sherratt, D. J. RecA bundles mediate homology pairing between distant sisters during DNA break repair. Nature 506, 249–253 (2014).

93 Dyck, E. V., Stasiak, A. Z., Stasiak, A. & West, S. C. Binding of double-strand breaks in DNA by human Rad52 protein. Nature 398, 728–731 (1999).

94 Fell, V. L. & Schild-Poulter, C. The Ku heterodimer: function in DNA repair and beyond. Mutation Research/Reviews in Mutation Research 763, 15–29 (2015).

95 Nimonkar, A. V., Sica, R. A. & Kowalczykowski, S. C. Rad52 promotes second-end DNA capture in double-stranded break repair to form complement-stabilized joint molecules. Proceedings of the National Academy of Sciences 106, 3077–3082 (2009).

96 McIlwraith, M. J. & West, S. C. DNA repair synthesis facilitates RAD52-mediated second-end capture during DSB repair. Molecular cell 29, 510–516 (2008).

