## Supplementary Information for "The human RAD52 complex undergoes phase separation and facilitates bundling and end-to-end tethering of RAD51 presynaptic filaments"

### Materials and Methods

#### Protein purification and fluorescence labeling

The human proteins 6xHis-tagged-RAD52[1], RAD51[2], and RPA[3] were expressed and purified to homogeneity using established protocols. The purified proteins were labeled via conjugating the N-terminal amine to an NHS ester conjugated fluorescent dyes (Lumiprobe) in a phosphate buffer (pH 7.0). The reaction was carried out at 10x dye to protein ratio for two hours at room temperature or overnight at 4 $^{o}$C. Ziba spin desalting columns (MWCO 7000) were used to separate the protein from the free dyes.

#### Phase separation analysis

RAD52 was prepared in mixtures containing 25 mM Tris-HCl (pH 7.5) and 30 mM NaCl in the presence of various concentrations of polymer crowder PEG8000 (M.W. 8 kDa, Fisher Scientific). The samples were mixed and immediately placed on a Lab-Tek 8 well chamber cover glass for subsequent imaging. For RAD52-ssDNA mixtures, RAD52 was mixed with the appropriate amounts of ssDNA dT40-Cy3 (IDT) at a protein concentration of 2.5 uM in a buffer containing 25 mM HCl and 15 mM NaCl. For RPA-RAD52 mixtures, we prepared solutions containing 1 uM of RAD52, 1 % (wt/vol) PEG8K, and variable RPA concentration in a buffer containing 25 mM Tris-HCl (pH 7.5) and 60 mM KCl. The salt identity and concentration were chosen partly based on the storage solution of the purified proteins since buffer exchange may lead to complex disassociation and reduced protein stability. All samples were placed on a Nikon eclipse Ti microscope equipped with a Nikon 100x oil immersion objective (Plan Apo 100x/1.4) and Andor iXon Ultra EMCCD camera. Each sample was imaged five times at different fields of view. For the multicomponent condensates preparation in Figure S4, we mixed RPA (1.2 µM + 300 nM RPA-MB543) and T30-ssDNA (with 200 nM Cy5-labeled 18mer DNA probe [GCCTCGCTGCCGTCGCCA]) in a buffer containing 25 mM Tris-HCl (pH 7.5), 30 mM NaCl, 40 mM KCl, and 1 % PEG. We then added RAD52 to a final concentration of 5 µM.

#### Recruitment experiments

Samples containing 5 µM concentration of RAD52 in 25 mM Tris-HCl (pH 7.5) and 30 mM NaCl in addition to nanomolar concentration of a fluorescently-labeled client (dT40-cy3, RPA-MB543, RAD51-AF 488, TERRA RNA-cy3) were added to the buffer prior to the addition of RAD52. The samples were placed into an 8-well Lab-Tek imaging chamber and loaded on a confocal microscope (Lumicks, Ctrap). Five images were taken for each sample for subsequent analysis of partition coefficients.

To calculate partition coefficients, we used ilastik pixel classification software[4] that relies on a neural network that can be trained manually by assigning pixel classification. We manually selected pixels that exist within condensates and pixels within the background and trained a model that uses features like intensity, intensity gradient, Laplacian, and other features to determine droplet pixels and background pixels. The analysis was then done on the entire image sets (5 images per sample) and outputs binary masks for the condensates. A custom-made python script is then used to identify condensates and compute their mean intensities and the background intensity. The partition coefficient for every condensate is then calculated and averaged for the entire sample. Error bars are estimated by calculating the standard deviation of partition coefficients.

#### RAD51 fibril sample preparation

Proteins and ssDNA were mixed in a 25 mM Tris-HCl (pH 7.5) and 100 mM NaCl buffer. The concentrations used for these experiments were 5 µM RAD51, 5 µM RAD52, 2.5 µM RPA, and 2.5 µM ssDNA. Different mixture combinations are shown in Figure 4 of the main text. We found that the order of addition is critical for these experiments. To induce fibril formation, ssDNA in added first to the buffer. Then, RAD51 is added and mixed with ssDNA before other protein components are added (such as RAD52 or RPA). All samples contained nanomolar concentrations of fluorescently-tagged proteins. The samples were loaded onto an objective-type total internal reflection fluorescence microscope (TIRF) and imaged in TIRF/HiLo mode. The

TIRF microscope is built on a Nikon eclipse Ti frame equipped with an Apo TIRF 60x/1.49 oil immersion objective and an Andor iXon Ultra EMCCD camera. For excitation, three laser lines are used (Coherent OBIS) with wavelengths 488, 561, and 640 nm.

**DNA PAINT super-resolution imaging**

RAD51 NPFs were prepared by mixing RAD51 and $\phi$X174 virion ssDNA at 2 μM RAD51 and 6 μM DNA (nucleotide concentration) in the reaction buffer, which contained 25 mM Hepes (pH 7.33), 30 mM KCl, 2 mM $CaCl_2$, 2 mM ATP, and 2 mM TCEP. The reaction was incubated for 15 minutes at 37 $^oC$. 0.1 μM RPA was added and the reaction was further incubated for 45 minutes. The sample is then deposited in an 8-well Lab-Tek chamber coated with poly(lysine) (M.W. ~1000-5000, P0879, Sigma-Aldrich) for 1 minute and then rinsed of with PBS. The sample is then blotted and left to dry in a fume hood for 20 minutes. Dehydration allows for better attachment of the complexes to the surface. We also checked the same samples without dehydration step and found no difference in the confirmation of filaments. Next, the sample is incubated with primary antibody against RAD51 (Rabbit, PC130, Sigma-Aldrich) at 4 $^oC$ overnight. For two color experiments, primary antibodies for RPA (Mouse, MA1-26418, ThermoFisher Scientific) or RAD52 (Mouse, MA5-31888, ThermoFisher Scientific) were also added. Next, the sample is washed three times with PBS followed by incubation of the oligo-conjugated secondary antibodies (Massive photonics, Fast sdAB 2-Plex kit) for 2 hours at room temperature. The sample is then washed three times with PBS. Imaging solution is added with 0.5-3 nM of imager strand concentration. For RAD51, the imagers were conjugated with ATTO-655 dye (Massive Photonics). For the RAD52 and RPA, the imagers were conjugated with Cy3B (Massive photonics). The sample was then loaded on a home-built objective-type TIRF microscope with an Olympus IX71 frame, a 100x/1.4 PlanSApo Olympus oil immersion objective and an Andor iXon EMCCD camera. The fluorescence illumination was achieved using a 500 mW 642 nm laser (MBP) and a 100 mW 568 nm laser (Coherent Saphire) that were used to excite ATTO655 and Cy3B, respectively. The microscope had a steering lens that controlled the angle of illumination that was set in TIRF mode. Movies were collected at an exposure time of 10 ms and for 10000-20000 frames. Image reconstruction, including fitting and rendering, was performed using the Picasso software from the Jungmann lab[5].

For strand invasion experiments, the homologous dsDNA ($\phi$X174 dsDNA RF1) was added and incubated for 30 minutes after the 1-hour-long assembly reaction had taken place. A similar procedure was performed for the RAD52-induced clustering with the following changes. RAD52 was added at 0.5 μM concentration and RPA was added at 1 μM at the start of the reaction. Then the reaction was incubated for 1 hr at 37 $^oC$ before imaging.

**Atomic Force Microscopy imaging**

Specification of High-Speed Atomic Force Microscope (HS-AFM)

A commercial Sample-Scanning High-Speed Atomic Force Microscope (SS-NEX Ando model) from RIBM (Research Institute of Biomolecule Metrology Co., Ltd.) was used for experiments involving RAD51 filaments and RAD51-RAD52 complexes. Tapping mode was employed to minimize interference with the deposited sample, and all deposited samples were captured in solution. Ultra-Short Cantilevers (USC-F1.2-k0.15-10), specifically designed for high-speed AFM with a resonance frequency of 1200 MHz, a spring constant of 0.15 N/m, and a length of 7 μm, were purchased from NanoAndMore and utilized in these experiments. A wide scanner was employed with scan speeds ranging from 0.05 to 1 frame per second, with the resolution set to 200 × 200 pixels.

AFM sample preparation

RAD51 filaments in the presence and absence of RAD52 were prepared and incubated in the same way as described earlier for DNA PAINT imaging. A freshly cleaved mica surface was coated with poly(lysine) (M.W. ~1000-5000, P0879, Sigma-Aldrich) at a concentration of 0.05 mg/ml. poly(lysine) was incubated for three minutes and washed off twice using distilled water. The RAD51 filament samples are diluted by a factor of 10 in the same buffer with ATP and then deposited on a freshly cleaved poly(lysine)-coated mica surface. After 10-minute incubation, the sample is washed with distilled water imaged with high-speed AFM microscope.

Image processing and analysis

HS-AFM images were viewed and analyzed using the software built by Prof. Toshio Ando's laboratory-built software, Kodec 4.4.7.39, with available source code[6]. Tilt and other image correction details are available in the literature[7]. Multiple frames were taken per structure and then projected with the median intensity value using Fiji-ImageJ software. The contrast was adjusted to enhance the structural features of the images.

**Quantification of phase separation from brightfield images**

To quantify the degree of phase separation from brightfield images, we used image analysis. Each sample was imaged five times at different fields of view within in the solution (away from the surface). Each bright field image was processed with a Farid filter, which is an image derivative filter that calculates intensity fluctuations in the XY plane. The resulting filtered image contains pixel values proportional to the gradient of intensity in the pixel. We then calculate the standard deviation of the pixel values in the filtered image. The filtered image is then passed through a logical filter with the condition (pixel value > 10 standard deviations), which will select the pixels containing high values of the image derivatives, which corresponds to sharp edges (i.e. edges of condensates in focus). The out-of-focus droplets are filtered out since the gradient of intensity will be lower than those that are in focus. This filtering of edges ensures that the background variations in intensity that could come from deposits on the surface and/or optical aberrations are not included in the calculations. Lastly, the number of selected pixels corresponding to the edges of droplets in focus is counted and divided by the area of the field of view, giving an estimate of the area fraction of droplet edges. This analysis is done for all five images and the resulting edge fractional area are average to give a quantitative indicator of the degree of phase separation. The errors are calculated as the standard deviation of the edge fractional areas of the five images. All the numbers are then normalized with the value obtained for a non-phase separating sample to remove the contribution of speckles within the microscope optics.

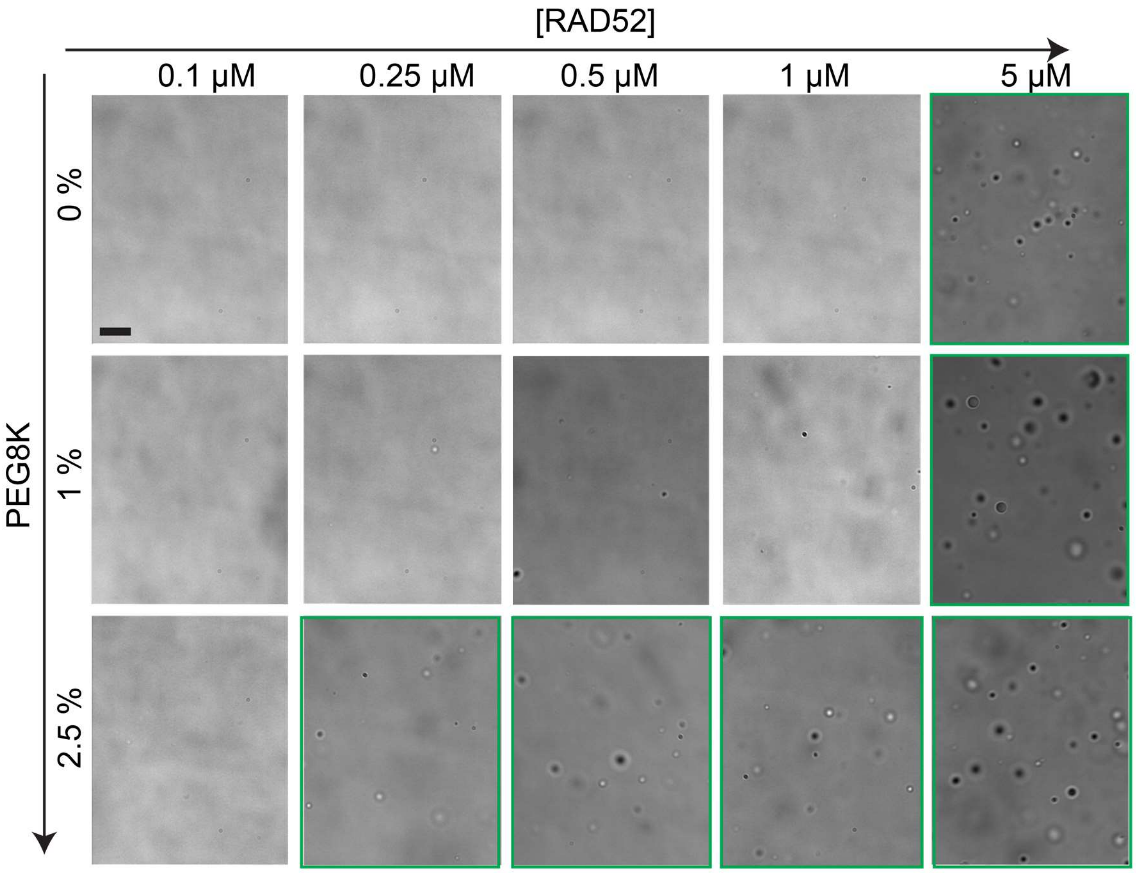


**Figure S1.** Bright-field images of RAD52 mixtures as a function of RAD52 concentration and PEG 8000 concentration (wt/vol). Scale bar is 10 µm. This is the complete set of conditions tested to compute the phase diagram in Figure 1f in the main text.

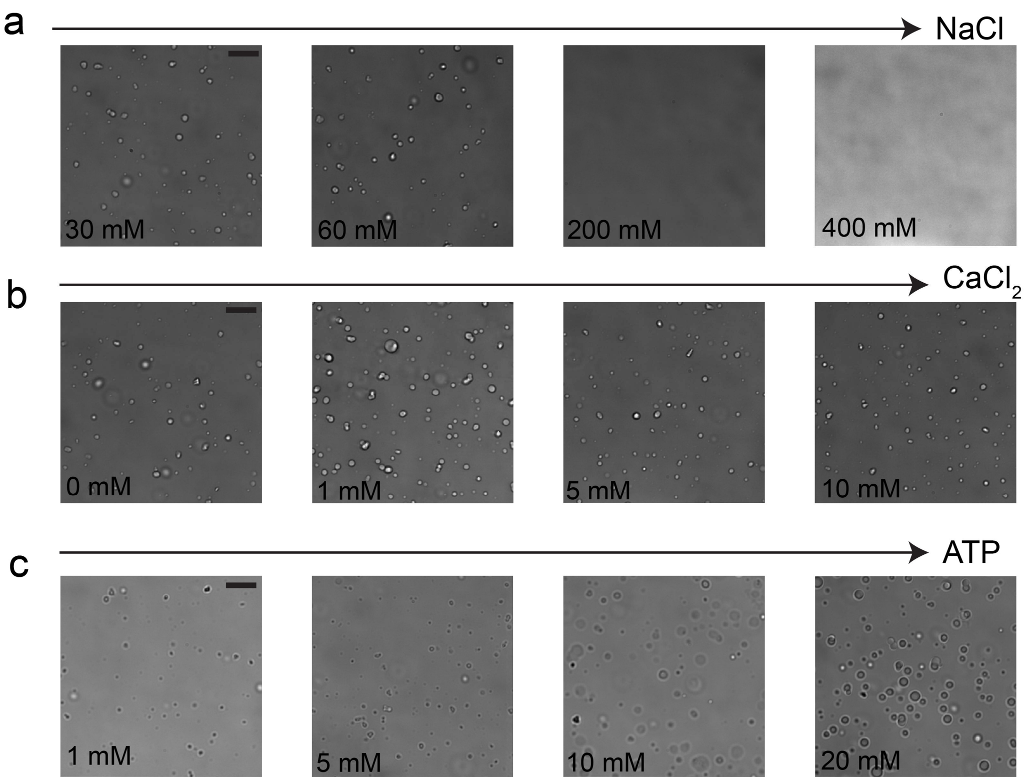


**Figure S2.** Bright-field images of RAD52 mixtures prepared at 5 μM RAD52 concentration and no crowder as a function of **(a)** NaCl, **(b)** $CaCl_2$, and **(c)** ATP concentrations. Scale bar is 10 μm. This is the complete set of conditions tested to compute the plots in Figure 1g-i in the main text.

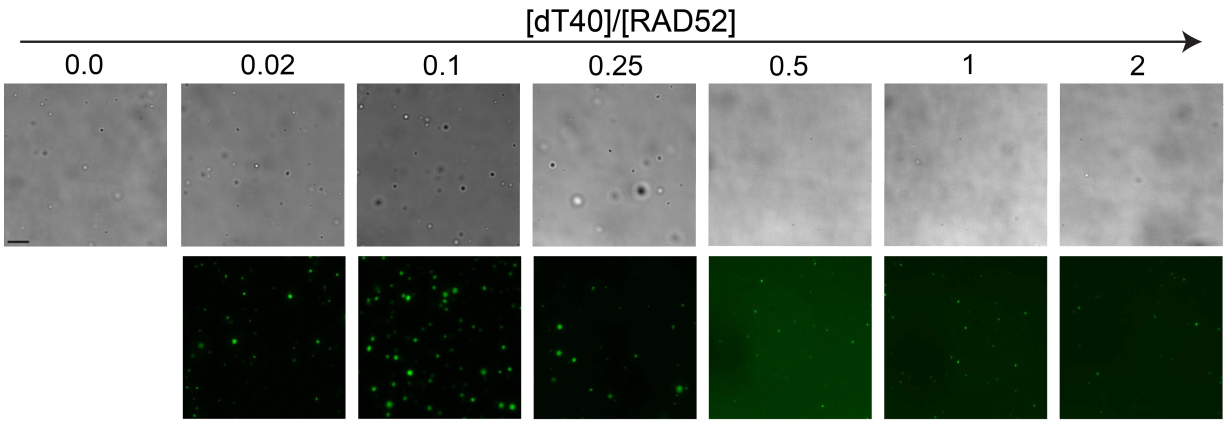


**Figure S3.** Bright-field (top) and fluorescence (bottom) images of RAD52-ssDNA mixtures prepared at 2.5 μM RAD52 concentration and variable ssDNA-to-protein ratio. Scale bar is 10 μm. This is the complete set of conditions tested that were used to compute the phase diagram plot in Figure 3b in the main text. dT40 ssDNA is conjugated to Cy3.

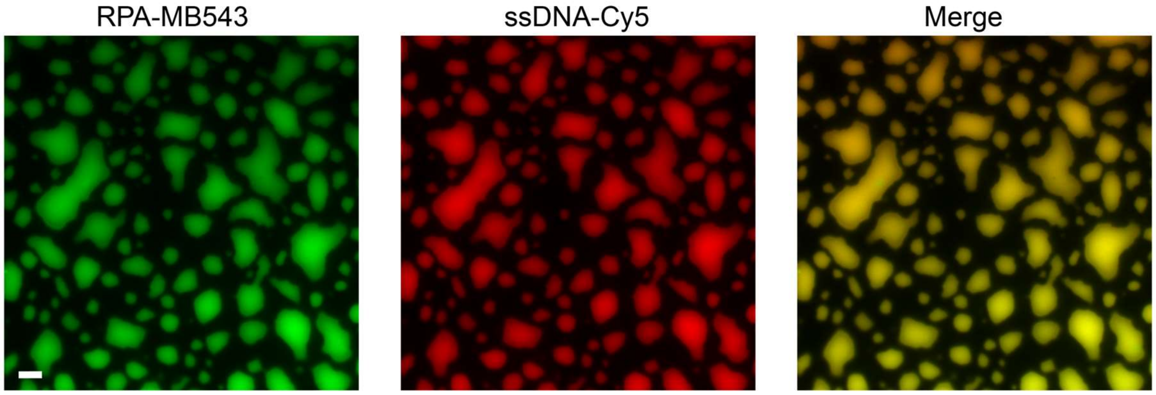


**Figure S4.** Multi-color fluorescence image of RAD52-dT40 condensates (with 200 nM Cy5-conjugated 18mer DNA probe) and RAD2-RPA (with 300 nM RPA-MB543) condensates mixed into one sample. Scale bar is 10 μm.

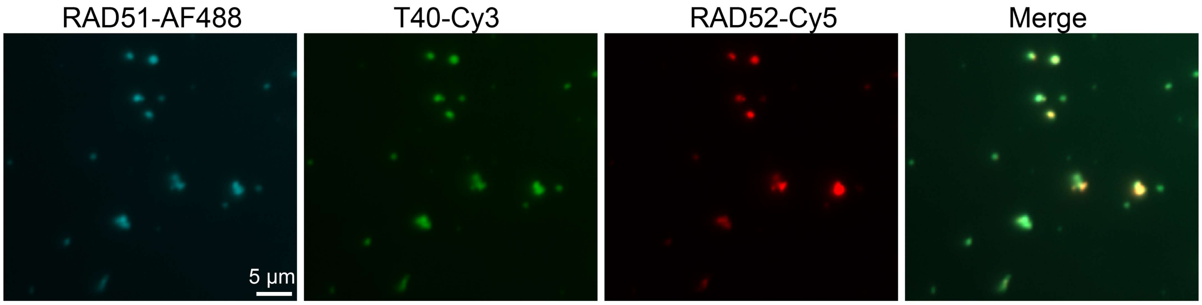


**Figure S5.** Multi-color fluorescence image of a RAD51-RAD52-ssDNA mixture. This sample is identical to the one shown in Figure 4d in the main text (no crowder condition), however, the two samples differ in the order of addition. In this sample, ssDNA is added first followed by RAD52 and then RAD51. In Figure 4d, ssDNA is added first followed by RAD51 and then RAD52.

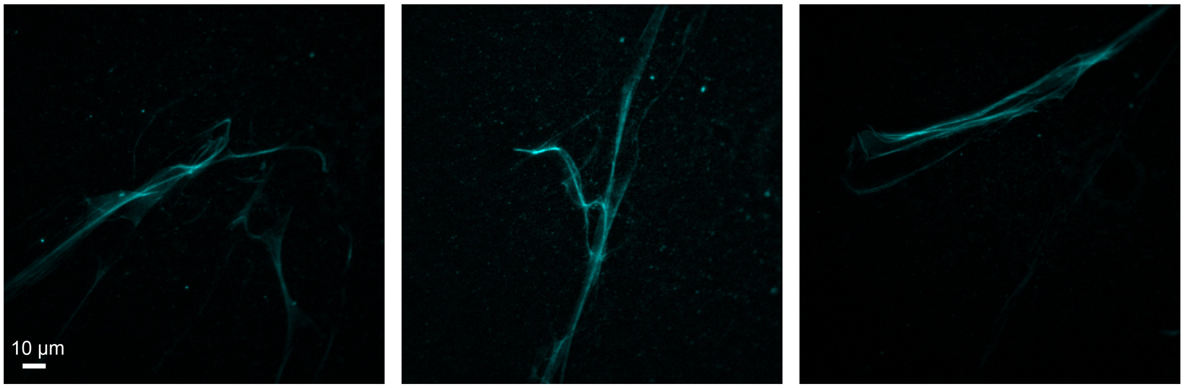


**Figure S6.** Fluorescence images of RAD51 (with 5% AF488-RAD51) fibers prepared by mixing RAD51 (5 µM) with ssDNA T30 (2.5 µM) and RAD52 (5 µM) in a buffer containing 25 mM Tris-HCl (pH 7.5), 100 mM NaCl, 2 mM ATP, and 2 mM $CaCl_2$.

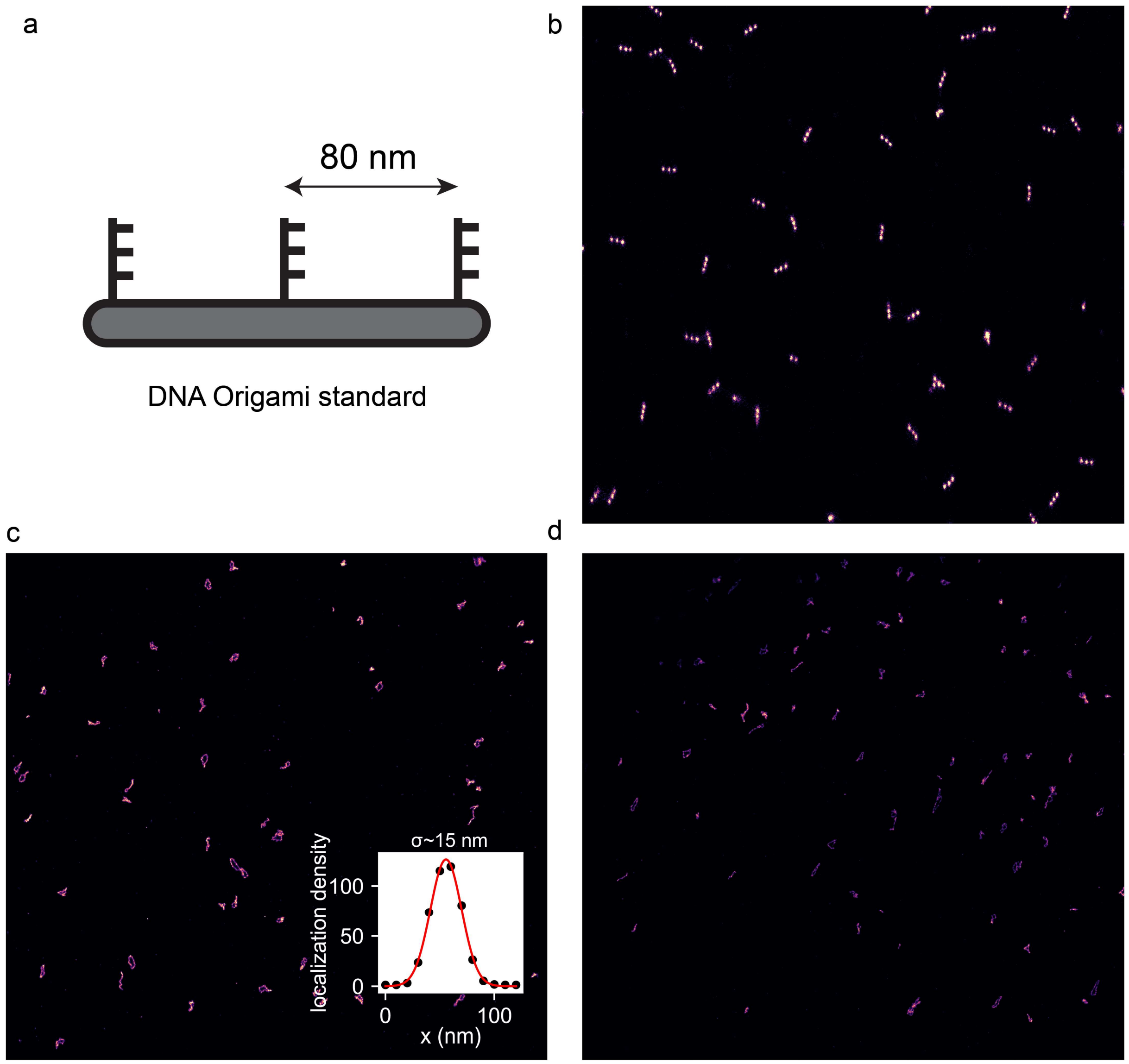


**Figure S7. (a)** Scheme describing a DNA origami nano-ruler (GATAquant) with probe strands located 80 nm apart that is used to judge the microscope resolution. **(b)** DNA paint image of the DNA nano-ruler **(c&d)** large view of the RAD51 NPF sample showing the circular and linear RAD51 NPFs formed on $\phi$X174 ssDNA. The standard deviation of the gaussian fit for a single protein localization is shown in the inset of **(c)** which gives an estimate of 15 nm resolution.

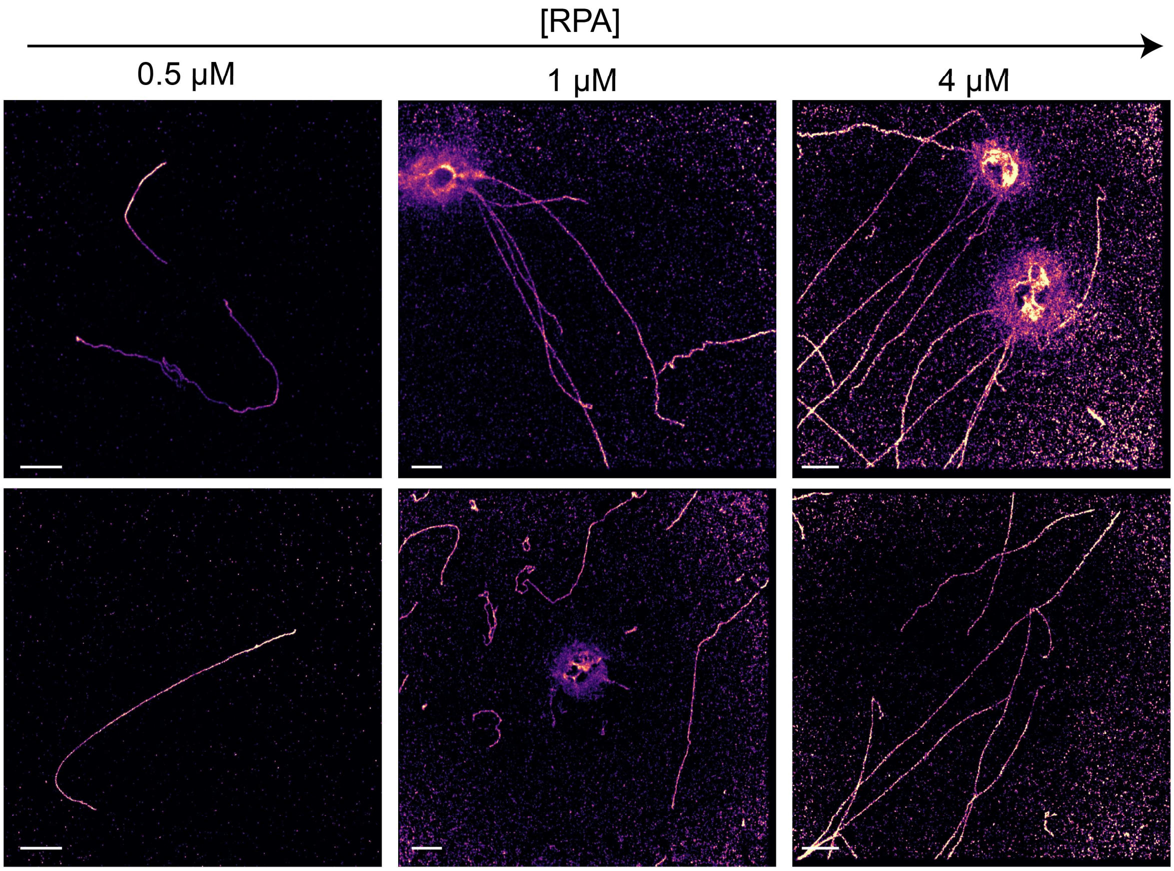


**Figure S8.** Super resolution DNA-PAINT images of RAD51 NPFs formed in the presence of 0.5 μM RAD52 and varying RPA concentration. All scale bars represent are 2 μm.

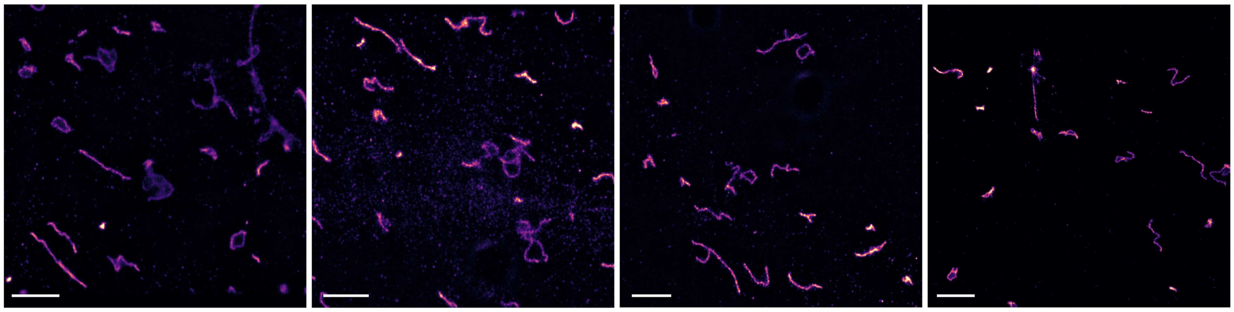

**Figure S9.** Super resolution DNA-PAINT images of RAD51 NPFs formed in the presence of 0.5 μM RAD52 on a $\phi$X174 dsDNA substrate. All scale bars represent are 2 μm.

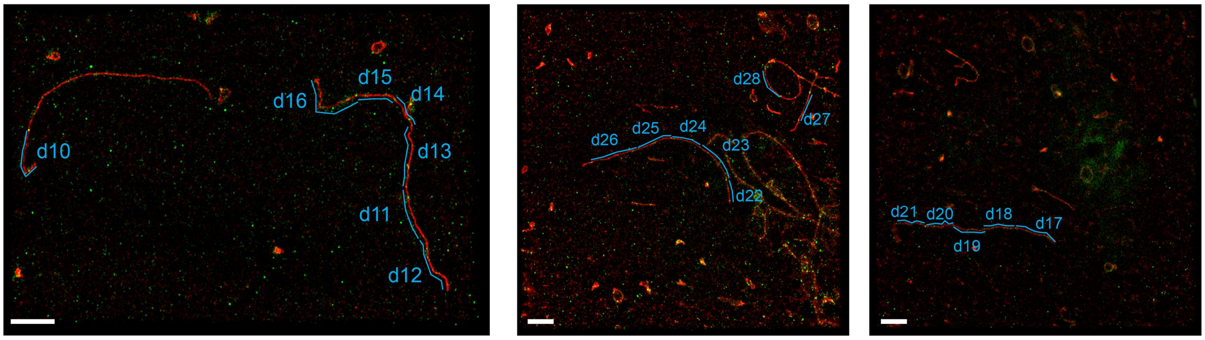


**Figure S10.** Two-color DNA-PAINT images of RAD51 NPFs formed in the presence of 0.5 μM RAD52 on a $\phi$X174 ssDNA substrate with staining for RAD52 (green) and RAD51 (red). All scale bars represent are 2 μm. The segmentation of the long filaments into shorter ones connected by RAD52 spots is used in Figure 6.

#### References


1 Grimme, J. M. *et al*. Human Rad52 binds and wraps single-stranded DNA and mediates annealing via two hRad52–ssDNA complexes. *Nucleic acids research* **38**, 2917-2930 (2010).

2 Subramanyam, S. & Spies, M. in *Methods in enzymology* Vol. 600 157-178 (Elsevier, 2018).

3 Henricksen, L. A., Umbricht, C. B. & Wold, M. S. Recombinant replication protein A: expression, complex formation, and functional characterization. *Journal of Biological Chemistry* **269**, 11121-11132 (1994).

4 Berg, S. *et al*. Ilastik: interactive machine learning for (bio) image analysis. *Nature methods* **16**, 1226-1232 (2019).

5 Schnitzbauer, J., Strauss, M. T., Schlichthaerle, T., Schueder, F. & Jungmann, R. Super-resolution microscopy with DNA-PAINT. *Nature protocols* **12**, 1198-1228 (2017).

6 Sakashita M, M. I., N Kodera, D Maruyama, H Watanabe, Y Moriguchi, and T Ando. Kodec4.4.7.39. (2013).

7 Ngo, K. X., Kodera, N., Katayama, E., Ando, T. & Uyeda, T. Q. Cofilin-induced unidirectional cooperative conformational changes in actin filaments revealed by high-speed atomic force microscopy. *elife* **4**, e04806 (2015).
